# Experimentally mimicking long term infections of *Magallana gigas* with the OsHV-1 virus reveals evolution through positive selection

**DOI:** 10.1101/2025.02.07.637049

**Authors:** Camille Pelletier, Nicole Faury, Mickaël Mege, Lionel Dégremont, Maelle Hattinguais, Jeremie Vidal-Dupiol, Germain Chevignon, Maude Jacquot, Benjamin Morga

## Abstract

Ostreid herpesvirus 1 (OsHV-1) poses a significant threat to the global oyster farming industry, causing substantial economic losses due to mortality outbreaks. While OsHV-1 primarily affects the Pacific oyster *Magallana gigas*, it has also been associated with mortality events in various other host species. Despite progress in understanding OsHV-1 epidemiology, important knowledge gaps remain regarding its evolutionary mechanisms and adaptation to host genetic backgrounds. This study uses experimental evolution and extensive genomic analysis to investigate the dynamics of OsHV-1 evolution in response to oyster host genetic variation. Our results show that genetic mutations, particularly transitions and transversions, play a key role in shaping viral populations, contributing to a trend toward genetic homogenization. Notably, stronger positive selection signals were observed in viral genomes isolated from oyster populations with higher susceptibility, suggesting adaptation of viral genotypes to specific host genetic backgrounds. These findings shed light on the complex evolutionary dynamics of OsHV-1 and its interactions with oyster hosts. Understanding how this virus adapts to host genetic diversity is crucial for developing strategies to mitigate its impact on the oyster farming industry and provides valuable insights into the broader mechanisms of viral evolution in response to host variation.

## 1. Introduction

Ostreid herpesvirus 1 (OsHV-1), a double-stranded DNA virus belonging to the *Malacoherpesviridae*, poses a significant threat to the global oyster farming industry by initiating a polymicrobial disease known as the Pacific Oyster Mortality Syndrome (POMS, de Lorgeril et al., 2018). In recent years, mortality outbreaks associated with POMS have caused substantial economic losses. Among several factors (*i.e.* temperature and food) that drive windows of oyster permissivity to disease, oyster age is a key determinant, with individuals being susceptible up to ∼18 months old (Azéma, Lamy, et al., 2017; Burioli et al., 2017; Nicolas et al., 1992; Renault et al., 1994). Meanwhile, oyster farmers have reported mortality associated with OsHV-1 mainly in spat, as survivors demonstrate genetic and epigenetic resistance upon subsequent exposure to viral particles (Evans et al., 2017; Gawra et al., 2023).

The first comprehensive characterization of the OsHV-1 genome was published in 2005 (Davison et al., 2005). The genome of OsHV-1 spans approximately 207 kilobase pairs (kb) in length and exhibits a distinctive genomic organization comprising unique regions (U_L_/U_S_: unique long or short) flanked by repeated regions (TR_L_/TR_S_: terminal repeat long or short and IR_L_/IR_S_: inverted repeat long or short). The architecture of the OsHV-1 genome can be summarized as TR_L_-U_L_-IR_L_-X-IR_S_-U_S_-TR_S_ (Davison et al., 2005). Advances in high-throughput sequencing have facilitated the genomic characterization of several OsHV-1 lineages, including OsHV-1 µVar (Burioli et al., 2017), OsHV-1 PT (Abbadi et al., 2018), and OsHV-1 SB (Xia et al., 2015). Currently, 28 complete genomes are available in public databases (Delmotte-Pelletier et al., 2022; Morga-Jacquot et al., 2021; Pelletier et al., 2025). These genomic datasets have revealed a spatiotemporal structuring of viral lineages in France (Delmotte-Pelletier et al., 2022) and globally (Morga-Jacquot et al., 2021), consistent with oyster farming practices. Furthermore, a recent study has highlighted a differentiation between lineages infecting *M. gigas* and *O. edulis*, with a greater viral diversity observed in *M. gigas* (Pelletier et al., 2025). However, despite significant advancements in understanding OsHV-1 epidemiology and evolution, several knowledge gaps persist regarding its evolutionary mechanisms and how it adapts to host genetic backgrounds. Indeed, substantial host genetic and epigenetic variation for resistance to OsHV-1 has been reported worldwide, enabling selective breeding programs to sustain oyster production (Camara et al., 2017; Dégremont et al., 2015b; Divilov et al., 2019; Gutierrez et al., 2020; Gawra et al., 2023; Valdivieso et al., 2025).

The evolution of viruses is driven by a range of processes, including mutation, genetic recombination, genetic drift, and natural selection. RNA viruses provide vivid examples of high mutation rates and rapid evolution. These viruses are known for their ability to accumulate genetic changes swiftly, resulting in substantial levels of viral polymorphism (Sanjuán, 2012; Sanjuán & Domingo-Calap, 2016, 2016). This genetic diversity empowers them to adapt to newly infected cellular environments and develop strategies to evade vaccines and antiviral drugs (Lauring, 2020). Although large dsDNA viruses exhibit lower mutation rates (*i.e.* between 10^-03^ and 10^-08^ nucleotide substitution/site/year, e.g. 5.9×10^-08^ for the Herpes simplex virus) compared to RNA viruses (*i.e.* between 10^-02^ and 10^-04^ nucleotide substitution/site/year) due to their use of high-fidelity proofreading polymerases (Sanjuán et al., 2010; Sanjuán, 2012; Sanjuán & Domingo-Calap, 2016), their genome stability is increasingly controversial as they sometimes display noteworthy genetic diversity (Sanjuán et al., 2016).

Viral evolutionary processes can be investigated through various approaches, including sequence analyses, metagenomics, phylogenetics, phylodynamics, and even Experimental Evolution (EE). This latter approach involves subjecting viral particles to controlled laboratory conditions, enabling the real-time observation and manipulation of evolutionary processes (Kawecki et al., 2012). The emergence and dynamics of mutations, immune escape variants, and evolutionary trade-offs can be assessed by monitoring viral populations under different selective pressures, such as antiviral drug treatments or interactions with immune systems (Kawecki et al., 2012).

While most studies using EE focus on easily replicable model organisms such as *Escherichia coli*, *Drosophila spp.*, *Saccharomyces cerevisiae* or various bacteriophages (Kawecki et al., 2012), recent research has expanded EE to diverse viruses, including Herpesviruses (Fuandila et al., 2022; Kuny et al., 2020).

For instance, *in vitro* EE conducted with populations of Herpes Simplex Virus type 1 (HSV-1) has revealed the accumulation of minor genetic variants and changes in population diversity. Although minor, these changes have been observed to lead to relatively rapid modifications in the viral population at the consensus level (Kuny et al., 2020). Another EE study, conducted on Koi herpesvirus (KHV), which infects the common carp *Cyprinus carpio*, has highlighted the key role of structural variations in driving rapid viral evolution *in vitro* (Fuandila et al., 2022). By driving genetic diversity, adaptation, and pathogenicity changes, these large-scale genomic alterations, including deletions, insertions, duplications, and inversions, can rapidly reshape viral genomes, influencing key functional regions (Fuandila et al., 2022).

In the absence of oyster cell lines for OsHV-1 propagation, we were forced to use *in vivo* techniques, which require high-throughput sequencing to efficiently and comprehensively analyze the evolution of viral populations. This however offered the advantage of studying this host pathogen interaction as a whole in an ecologically relevant context.

In the context of OsHV-1, our understanding of the evolutionary processes contributing to the diversity of the viral population remains limited. The primary objective of the present study was to gain insights into the evolutionary mechanisms driving the adaptation and diversification of OsHV-1 lineages based on oyster susceptibility to infection. To achieve this goal, we used an EE approach spanning up to 28 passages under controlled conditions, combined with a comprehensive sequencing analysis of 96 OsHV-1 genomes.

## 2. Material and methods

### 2.1. Oysters production

To explore the impact of oyster genetic backgrounds on the evolutionary dynamics of OsHV-1, we generated three oyster populations with different levels of resistance to OsHV-1 infection. In December 2021, wild oysters were collected from two sites in Charente-Maritime (France) based on their proximity to oyster farms and history of viral exposure.

The first site, La Floride (Marennes-Oléron Bay, Charente-Maritime, Lat: 45.803; Long.: -1.153) is surrounded by oyster farms, densely populated by wild oyster beds and is exposed to OsHV-1 outbreaks annually (Dégremont et al., 2019). Oysters from this site have shown high resistance to POMS (Valdivieso et al., 2025).

The second site, Chaucre (Charente-Maritime, Lat: 45.982; Long: -1.396), is located on the west coast of Oléron Island in a low-density, non-farming area. Wild oysters collected from Chaucre are less frequently exposed to OsHV-1 µVar than oysters from densely populated areas and are therefore considered moderately susceptible to OsHV-1 µVar infection.

In addition, a control oyster population used in previous studies was included (Gawra et al., 2023; Valdivieso et al., 2025). These oysters were initially sampled in 2008 prior to the emergence of OsHV-1 µVar, and have since been bred for six generations using oysters from second-passage individuals never exposed to OsHV-1 µVar. This lineage has been maintained in a secure facility with UV-treated seawater (40 mJ/cm²), and is considered naive to OsHV-1 µVar since 2009. It has been described as highly susceptible to OsHV-1 µVar, with mortality rates exceeding 80% (Valdivieso et al., 2025).

The three oyster populations (La Floride, Chaucre, and the naïve control) were reared separately at the Ifremer hatchery in La Tremblade (December 2021) to prevent cross-contamination with pathogens such as OsHV-1 and *Vibrio aestuarianus* (Dégremont et al., 2005). Seawater temperature was gradually increased from 11°C to 20°C and oysters were fed with a cultured phytoplankton mix (*Isochrysis galbana, Tetraselmis suecica* and *Skeletonema costatum*) to induce gametogenesis (Dégremont et al., 2005).

In March 2022, 30 oysters per population were sexed, and gametes were collected via gonad stripping. Female gametes were pooled and filtered through 20-μm and 100-μm meshes, then fertilized with sperm from individual males. After 10 minutes, all fertilized eggs were pooled and transferred to 30 L tanks to minimize sperm competition and maximize effective population size (Boudry et al., 2002). Larval and spat culture followed established protocols (Dégremont et al., 2005, 2007). Progenies were reared in UV-treated seawater until experimental OsHV-1 infections and designated as FA, NFA, and C, corresponding to farming area (high density with the typical oyster bed of 100 individuals/m² and annual POMS reported), non-farming area (low-density beds, less than 20 individuals/m² and no POMS reported), and control populations (naïve laboratory-reared population, highly sensitive to OsHV-1 infection).

Before the experiment, spat were acclimated in 120 L tanks with a continuous flow of UV-treated seawater heated to 19°C. The seawater was supplemented with a phytoplankton mixture containing *I. galbana, T. suecica* and *S. costatum*. The acclimation period lasted at least 2 weeks to ensure optimal growth and feeding conditions, critical for efficient OsHV-1 replication and observable oyster mortality (Azéma, Maurouard, et al., 2017).

### 2.2. Virion suspension preparation

To generate a diversified viral inoculum for infecting all oyster populations, nine OsHV-1 suspensions were produced between 2020 and 2021 (Pelletier et al., 2026). These suspensions were obtained from spat oysters naturally infected during field monitoring of OsHV-1 µVar outbreaks in various areas in France. Briefly, approximately 1000 specific-pathogen-free (SPF) spat oysters, reared at the Ifremer experimental facility in Argenton (Brittany, France), following a standardized protocol (Petton et al., 2015), were deployed in four oyster farming areas during active disease outbreaks (Figure 1). These areas included: *i)* Brest harbor (BR) (used to produce two virion suspensions) (Logonna-Daoulas, lat.: 48.335 long.: -4.318), *ii)* Marennes-Oléron Bay (MO) (used to produce four virion suspensions) (La Floride, lat.: 45.803 and long.: -1.153), *iii)* Arcachon Basin (ARC) (used to produce two virion suspensions) (Grahude, lat.: 44.653 and long.: -1.073), *iv)* Leucate lagoon (LEU) (used to produce one virion suspension) (Leucate, lat.: 43.379 and long.: 3.571). Seven days after deployment of SPF spat, once mortalities began, oysters were retrieved and kept at 20°C for 7 days in the lab. During this period, moribund oysters were sampled daily and stored at -80°C. A total of 126 samples were selected for whole-genome sequencing, including *i)* 10 samples collected in 2020 and 15 in 2021 from Brest harbor, *ii)* 20 samples in 2020 and 44 in 2021 from Marennes-Oléron Bay, *iii)* 10 samples in 2020 and 15 in 2021 from Arcachon Basin, and *iv)* 12 samples collected in 2021 from Leucate lagoon (Pelletier et al. 2026). In total, nine virion suspensions were prepared following a standardized protocol (Schikorski et al., 2011) (Figure 1, Table S1).

**Figure 1:**
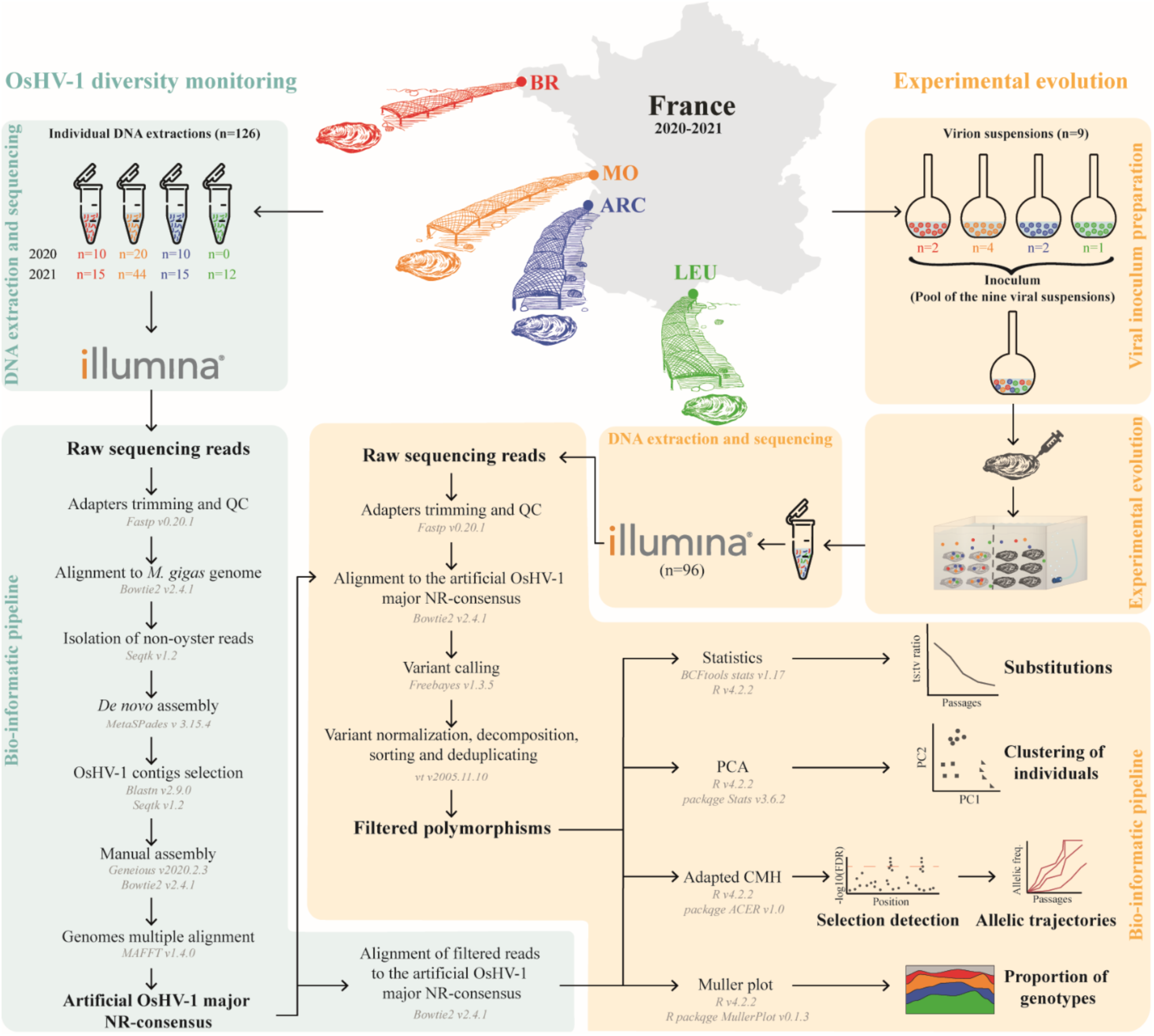
Overview of the bioinformatic workflow used to monitor OsHV-1 genomic diversity and to detect selection patterns during experimental evolution. The central panel shows the sampling map with the four French oyster-farming areas involved in the study between 2020 and 2021: BR (Bittany), MO (Marennes-Oléron), ARC (Arcachon), and LEU (Leucate). The number of samples processed in each region is indicated for both 2020 and 2021, with color-coded oyster illustrations representing each site. The left section (blue panels) describes the analytical pipeline applied to naturally infected oysters collected between 2020 and 2021 from four farming areas in France (Pelletier et al. 2026). Virion DNA was individually extracted from mantle tissue of moribund oysters and sequenced using Illumina technology. Raw reads were assembled *de novo* into OsHV-1 NR-genomes using a dedicated bioinformatic pipeline previously developed for OsHV-1 genomic analysis (Dotto-Maurel-Pelletier et al., 2019). The right section (orange panels) illustrates the experimental evolution design and associated bioinformatic analyses. Nine virion suspensions were prepared from moribund oysters collected in the field, then pooled in equimolar proportions to create a standardized inoculum. This inoculum was used to infect oysters in a controlled evolution experiment (see Figure 2 for details). DNA extracted from infected oysters at successive time points was sequenced using Illumina technology. A specific bioinformatic pipeline was developed to detect and characterize signatures of selection.

### 2.3. Inoculum virion load quantification and virion suspension mix setup

To prepare an equimolar viral inoculum for injection into oysters, viral particles of each of the nine virion suspensions were quantified. DNA was extracted from 100 µL of each virion suspension using the MagAttract^®^ HMW DNA kit according to the manufacturer’s instructions. DNA purity and concentration were measured using a Nano-Drop ND-1000 spectrophotometer (Thermo Scientific) and Qubit^®^ dsDNA BR assay kits (Molecular Probes Life Technologies), respectively. Viral DNA copy number was then quantified by quantitative PCR using a Mx3005 P Thermocycler (Agilent) (Pepin et al. 2008). Each reaction contained 1 µL of Milli-Q water, 2 µL of each primer at 5.5 µM, OsHVDP For (forward) 5′-ATTGATGATGTGGATAATCTGTG-3′ and OsHVDP Rev (reverse) 5′-GGTAAATACCATTGGTCTTGTTCC-3′, 10 µL of Brilliant III Ultra-Fast SYBR® Green PCR Master Mix (Agilent), and finally 25 ng of DNA sample from infected oysters or Milli-Q water (non-template control) in a total volume of 20 µL. The qPCR program consisted of 3 min at 95 °C followed by 40 cycles of amplification at 95 °C for 5s and 60 °C for 20s. Results were expressed as the log-transformed copy number of viral DNA per microliter of seawater (cp/µL).

The nine virion suspensions were pooled to create the initial virion inoculum (Table S1). To ensure equal contribution from each suspension, the pool was adjusted to obtain an equimolar concentration of virion DNA (Table S1).

### 2.4. Experimental evolution design

The EE was conducted under controlled conditions in a Type 2 laboratory at Ifremer’s La Tremblade facility. To initiate the experimental infections, 50 oysters from each population were first myorelaxed using hexahydrate MgCl_2_ (50 g/L) (Suquet et al., 2009). Each oyster was then individually injected with 100 μL of pooled inoculum into the abductor muscle using a 26-gauge needle attached to a multi-dispensing hand pipette (Figure 2, step 1). At the same time, 10 oysters per population were myorelaxed and injected with artificial seawater to serve as control. The experiment was conducted in duplicate (*i.e*. two tanks per condition), with oysters placed in 5-L plastic tanks filled with seawater at 20°C and continuously supplied with a phytoplankton mixture (*Isochrysis galbana, Tetraselmis suecica* and *Skeletonema costatum*). Mortality was monitored daily for seven days.

**Figure 2:**
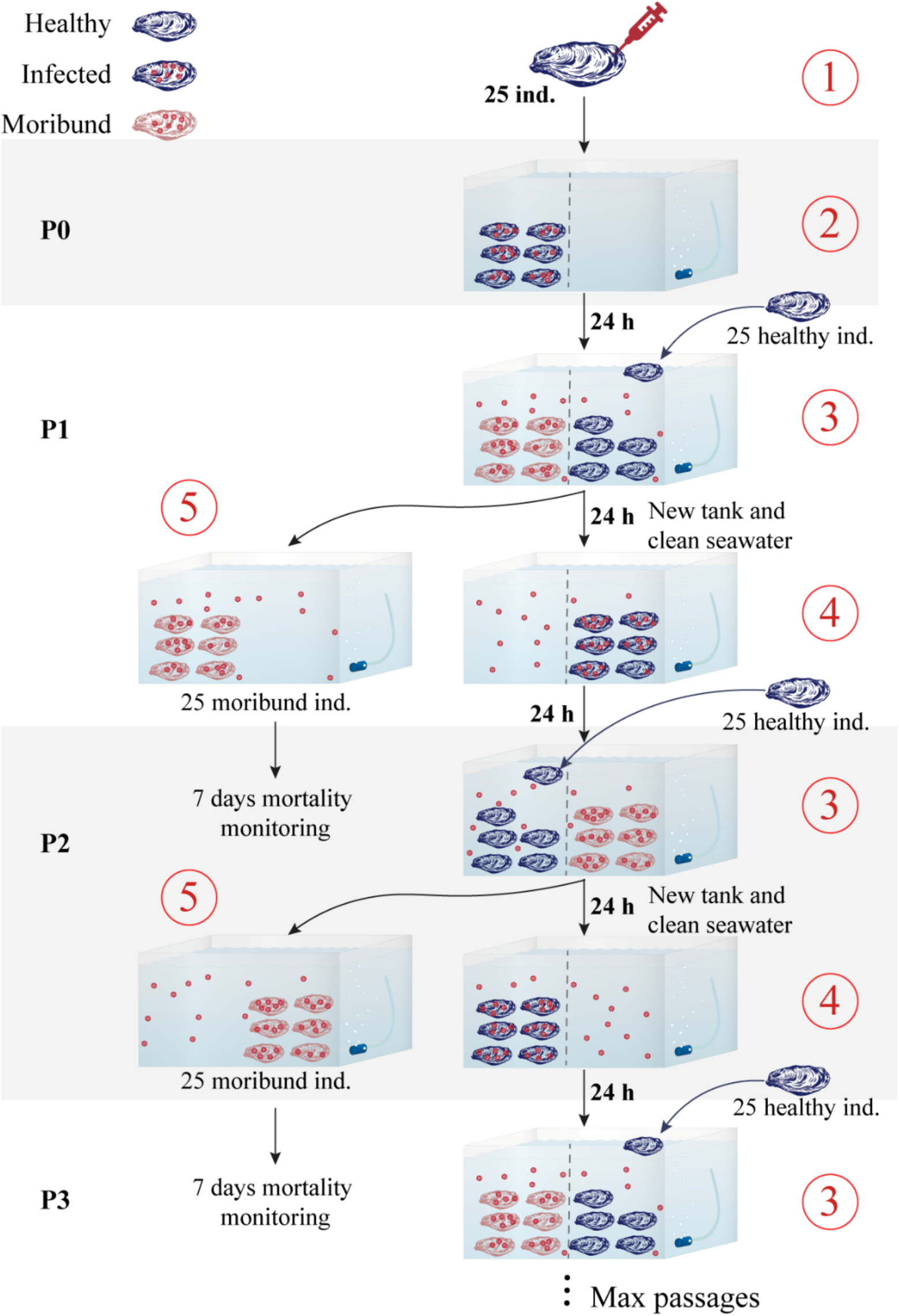
Experimental evolution design for one oyster population. The experimental evolution was conducted independently on three oyster populations using a serial cohabitation infection design. Healthy oysters are shown in dark blue, infected individuals in dark blue with red viral particles, and moribund oysters in light red with red viral particles. Experiments were performed in duplicate for each oyster population. At passage P0, 25 healthy oysters were injected intramuscularly with a standardized viral inoculum to initiate infection (step 1). After 24 hours, the infected oysters were transferred to tanks containing clean seawater (step 2). A new batch of 25 healthy oysters was then introduced to allow cohabitation and passive exposure to viral particles released into the water (step 3). After 24 hours, exposed oysters were moved to fresh tanks with pathogen-free seawater (step 4) to allow the progression of infection under controlled conditions. Moribund individuals were monitored over a 7-day period (step 5), and the 25 newly infected oysters were used to initiate the next passage (P1). This serial transfer process was repeated for each passage until the designated endpoint, with each round involving exposure, infection, and mortality monitoring.

Following this initial infection, nine tanks were prepared: two containing infected oysters and one containing ASW-injected control oysters for each oyster population. These tanks served as the starting point, referred as “Passage 0” (Figure 2, step 2). To initiate the first transmission passage (P1) through cohabitation, 25 healthy oysters from each population were introduced into the corresponding infected tanks 24 hours after donor injection (Figure 2, step 3). For instance, 25 FA oysters were added to each FA-infected tank, and so on. After 24 hours of cohabitation, these healthy oysters were transferred to fresh, pathogen-free seawater to allow virion excretion and initiate the next infection passage (Figure 2, step 4). This procedure was repeated to produce a second passage, following the same protocol. During each cohabitation period, donor oysters were monitored for seven days, and all moribund oysters were collected for DNA extraction (Figure 2, step 5). In total, we completed 13, 18, and 28 passages for the FA, C, and NFA oyster populations, respectively. Mortality gradually declined across all experimental populations, indicating the cessation of active virion transmission and marking the end of the experimental infection.

### 2.5. DNA extraction and sequencing

DNA extraction was performed on oysters collected during mortality monitoring (Figure 2, step 5) at specific infection passages: P0, P5, P10, and P13 for the FA population, at passages P0, P5, P10, P15, and P18 for the C population, and at passages P0, P5, P10, P15, P20, P24, and P28 for the NFA population. Sampling every five passages was chosen to limit the number of samples processed while ensuring consistent infection dynamics. This approach explains the discrete distribution of viral genome copy numbers and confirms that virion loads remained relatively stable across passages.

For each oyster population, infection passages, and replicate, seven samples of 35 mg of mantle tissue were collected for DNA extraction, which were performed using the MagAttract® HMW DNA kit (Qiagen) following manufacturer’s instructions. Sample selection was based on mortality dynamics during the experimental exposure. Individuals were preferentially collected on the day corresponding to the highest observed mortality, in order to characterize viral genome loads during periods of maximal disease impact. When sampling at the exact mortality peak was not possible, individuals were selected on days immediately before or after this peak, based on observed mortality patterns and the availability of intact animals, as advanced tissue decomposition in dead oysters sometimes precluded reliable sampling. Samples were collected from each replicate accordingly.

DNA purity and concentration were measured using a Nano-Drop ND-1000 spectrophotometer (Thermo Scientific) and Qubit® dsDNA BR assay kits (Molecular Probes Life Technologies), respectively. Viral genome copy number was quantified by quantitative PCR using a Mx3005 P Thermocycler (Agilent), as previously described. From each oyster population and replicate, three individuals were selected according to their OsHV-1 viral genome copy number and DNA concentration, to ensure sampling of oysters with representative viral genome copy number. In total, 96 samples were sequenced by DNA-seq Illumina by IntegraGen SA (Evry, France). PCR-free libraries were prepared using the Twist library Preparation Enzymatic Fragmentation (EF) Kit 1.0 (Twist Bioscience) according to the supplier’s recommendations. Briefly, after quantification of double-stranded genomic DNA, 400 ng of DNA were enzymatically fragmented to generate inserts of approximately 400 bp. Following fragmentation, libraries underwent end-repair, A-tailing, ligation with unique dual-index (UDI) Illumina adapters. Libraries were then purified and size-selected using SPRI beads, quantified by qPCR and sequenced on the Illumina NovaSeq platform to generate paired-end 150 bp reads.

### 2.6. Statistical analysis of mortality and OsHV-1 virion load across passages and oyster populations

Mortality and virion load data were analyzed using R v4.2.2. Raw data were graphically represented using ggplot2 v3.5.2. Differences in mortality and viral load among generations and populations were assessed using an ANOVA analysis. Linear mixed-effects models were fitted using the function lmer from the lmerTest package v3.2-0 (Kuznetsova et al., 2017) with replicates included as a random effect. The significance of fixed effects was assessed on the fitted mixed models using anova function from the stats package v4.5.1 (R Core Team, 2019).

To assess the consistency of mortality dynamics among biological replicates, a global covariance structure was estimated by computing pairwise Pearson correlation coefficients between replicate time series. A correlation matrix was generated using pairwise complete observations to account for missing values. This matrix quantifies the strength and direction of co-variation in mortality rates among replicates across passages and was visualized as a heatmap to facilitate interpretation of concordance patterns between experimental lines.

### 2.7. Assembly of a global consensus for each virion suspension

To reconstruct the overall genetic composition of the viral inocula, we assembled 126 de novo OsHV-1 µVar NR-genomes from infected oyster tissues corresponding to the batches used for virion suspension preparation (Figure 1, part blue, Table S2). DNA was extracted, quantified, and sequenced as previously described (see “DNA extraction and sequencing” section). DNA libraries were prepared using the Shotgun PCR-free library preparation kit (Lucigen) and sequenced on an Illumina NovaSeq^TM^ 6000 system (paired-ends, 150 bp reads) by Genome Quebec Innovation Center (McGill University, Montreal, Canada).

Raw sequencing data are publicly available in the NCBI SRA database under Bioproject PRJNA1216400 with accession numbers SAMN46433918 to SAMN46434013 to ensure data accessibility and reproducibility (Table S2).

*De novo* OsHV-1 NR-genome assemblies were generated from sequencing reads as previously described using a custom bioinformatic pipeline (see Dotto-Maurel et al., 2022 for details on the bioinformatic pipeline). Briefly, for each library, sequencing reads were quality-filtered (PHRED score > 31) and trimmed using Fastp v0.20.1 (with parameter -q 31; Chen et al., 2018). Filtered reads were mapped to the *M. gigas* reference genome (accession number: GCF_902806645.1; Peñaloza et al., 2021) using Bowtie2 v2.4.1 (parameter --local; Langmead & Salzberg, 2012) to remove host sequences. The remaining reads were assembled *de novo* using metaSPAdes v3.15.4 (parameter --meta; Bankevich et al., 2012). OsHV-1 contigs were identified with BLASTn v2.12.0 (options -max_target_seqs 1, -evalue “1e-10”; Altschul et al., 1990), manually re-ordered in Geneious v2022.2.2 based on read overlaps, and reconstructed as NR-genomes to remove redundant inverted repeats. Assembly accuracy and coverage uniformity were validated by dotplot similarity and read realignment with Bowtie2.

To summarize the genetic diversity of the viral suspensions, the 126 assembled NR-genomes were aligned using MAFFT v1.4.0 (Katoh et al., 2002, Figure 1). In this manuscript, the resulting consensus is referred as “artificial OsHV-1 major NR-consensus”.

### 2.8. Variant calling

To assess the genomic variability among individuals, intra-individual Single Nucleotide Variants (iSNVs) - defined as sites displaying at least two alleles with frequencies above 10% in at least one sample – were identified for each sequencing library based on the “artificial OsHV-1 major NR-consensus” previously generated. Variant calling was conducted using a modified version of a previously described pipeline for detecting polymorphisms (Figure 1, see Delmotte-Pelletier et al., 2022 for details of the bioinformatic workflow). Briefly, reads were filtered using Fastp v0.20.1 (Chen et al., 2018), mapped to the “artificial OsHV-1 major NR-consensus” using Bowtie2 v2.4.1 (Langmead & Salzberg, 2012) and variants were called using Freebayes v1.3.5 (Garrison & Marth, 2012), with the following settings: *-use-mapping-quality, -min-repeat-entropy 1, -haplotype-length 0, -min-alternate-count 5, -pooled-continuous, -hwe-priors-off, -allele-balance-priors-off*. The resulting variant calling outputs were normalized, decomposed, sorted and deduplicated using vt v2005.11.10 (Tan et al., 2015). Only libraries with a minimum mean genome-wide coverage of 110× were retained for downstream analyses (Delmotte et al., 2022; Pelletier et al., 2025). Variant calling focused on SNVs (transitions and transversions), while indels (16 insertions and 190 deletions) were excluded from downstream analyses because they were few, mostly located in complex regions such as homopolymeric G or C stretches, and could not be reliably analyzed with post-hoc methods such as Cochran–Mantel– Haenszel tests or allelic frequency plots. iSNVs were included in the analysis only if their frequency was ≥10% in individuals. Detected iSNVs were then mapped to their respective consensus genomes and aligned using MAFFT v1.4.0 (Katoh et al., 2002) implemented in Geneious Prime to re-coordinate all iSNVs positions across samples. Each re-coordinated iSNV was assigned a unique identifier based on *(i*) its position within the multiple alignment, (*ii*) the reference allele (REF) and *(iii*) the alternative allele (ALT) (*i.e.* position_REF >ALT) following the approach described by Delmotte-Pelletier et al. (2022).

### 2.9. Analysis of substitutions types and clustering of individuals based on iSNVs

To obtain detailed information on the number of transitions (*i.e.* substitution of a purine by another purine or a pyrimidine by another pyrimidine) and transversions (*i.e.* substitution of a purine by a pyrimidine or a pyrimidine by a purine) within OsHV-1 genome across passages and among host oyster populations, iSNVs detected in all libraries were analyzed using BCFtools stats v1.17 (Figure 1, Danecek et al., 2021). In the absence of a true ancestral genome, substitution directionality cannot be inferred with certainty, consequently, complementary substitutions (*e.g.* A>G and G>A) were collapsed into base pair substitution classes (*e.g.* A|G). In addition, iSNVs were analyzed separately depending on whether they were identified as ancestral or *de novo*, allowing distinction between newly emerging iSNVs and those inherited from previous viral passages. Ancestral status was determined at P0 by counting the number of individuals across both replicates in which the alternative allele was detected (allele frequency > 0). iSNVs present in more than one individual were classified as ancestral, whereas those detected in zero or one individual were considered *de novo*. For substitution pattern analyses, iSNVs counts were first computed independently for each library by recording the number of genomic positions showing a non-zero allele frequency for each substitution type. Identical iSNVs detected in multiple individuals for each replicate were subsequently averaged across individuals and replicates within each oyster population and passage. Substitution counts were then averaged per passage and per oyster population and visualized using R v4.2.2 and the ggplot2 package v3.4.0 (Wickham, 2009).

The ts:tv ratios were calculated to compare substitution patterns among samples, and a baseline ts:tv ratio of 0.5 was used to represent the expected value under random sequencing errors (Zook et al., 2014). Nucleotide diversity (π) was calculated from allele frequency data. For each iSNV and sample, the alternative allele frequency was extracted and π was computed using the classical biallelic formulation (Nelson & Hughes, 2015)

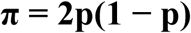

where p corresponds to the alternative allele frequency. This metric represents the expected heterozygosity at a given site, i.e., the probability that two randomly sampled genomes differ at that position. Site-specific π values were then averaged across all positions to obtain a mean intra-individual nucleotide diversity. Mean π and standard deviation were subsequently calculated per family and per passage, providing a summary of within-population genetic diversity dynamics over time.

### 2.10. Exploring sample clustering and iSNV data quality assessment through Principal Component Analysis

In the context of EE studies, Principal Component Analysis (PCA) was employed to assess sample clustering and evaluate the quality of intra-individual Single Nucleotide Variants (iSNVs) data, allowing detection of underlying population structure and potential data anomalies (Patterson et al., 2006).

Based on iSNVs frequencies, individuals were projected into a two-dimensional space to assess sample clustering. PCA was performed using the pcrcomp function from the factoextra v1.0.7 package (Kassambara & Mundt, 2020) in R v4.2.2 (Figure 1) based on a vector of 1,042 positions to capture intra-individual variation. Cluster groupings were visualized using ellipses drawn with stat_ellipse() in ggplot2. This analysis incorporated information on oyster populations, infection passages and replicate identity.

### 2.11. Selection detection

Selection patterns were detected using a SNV-based approach. Since the experiment was conducted in duplicate for each oyster population, selected iSNVs could appear in both replicates or only in specific ones. Selection of particular OsHV-1 lineages occurred independently within each oyster population, and consequently, all analyses were performed separately for each population. To detect iSNVs showing significant shifts in allele frequency (AF), we compared the initial passages (P0) with the evolved passages (up to P13, P18, and P28 for FA, C and NFA oyster populations, respectively). For this purpose, an adapted Cochran-Mantel-Haenszel (CMH) test was applied (Figure 1) using the ACER v1.0 package (Barghi et al., 2020; Spitzer et al., 2020) in R. The CMH test was parameterized using matrices of individual iSNV allele frequencies and individual sequencing coverages, together with the effective population size (Ne), the number of passages included, the number of replicates, and the pool size corresponding to the three individuals sequenced per replicate. This framework explicitly accounts for pooled individuals and replicate structure, and compute p-values for each iSNV, reflecting frequency changes over time while accounting for genetic drift and sampling variance (Spitzer et al., 2020). As an intrinsic property of the CMH approach, a single p-value was obtained per iSNV, population, and passage, reflecting parallel frequency changes across pooled individuals and replicates rather than collapsing data arbitrarily.

P-values were then adjusted for multiple testing using the false discovery rate (FDR) method, with the p.adjust function from the stats v4.2.2 package (R Development Core Team, 2005). Adjusted p-values were visualized on the OsHV-1 NR-genome using Manhattan plots generated with the ggplot2 v3.4.0 package (Wickham, 2009). iSNVs were considered under selection if adjusted p-values exceeded the CMH threshold corresponding to a 5% FDR. Finally, allele frequency trajectories across passages were plotted for each replicate using the ggplot2 v3.4.0 package (Wickham, 2009), based on adjusted p-values and frequency data, with allele frequencies calculated from pooled individuals within each replicate.

Local haplotype structure was inferred from iSNV frequency data by grouping spatially proximate polymorphic sites along the OsHV-1 genome. iSNV positions were first ordered according to their genomic coordinates, and contiguous blocks were defined using a sliding distance criterion: consecutive iSNVs separated by ≤300 bp were assigned to the same block, whereas larger gaps initiated a new block. This procedure partitions the genome into clusters of closely spaced variants that are likely to be physically linked and therefore potentially co-inherited. The resulting table provides a synthetic overview of local haplotype organization across the genome, highlighting clusters of mutations that may reflect linked evolutionary events.

### 2.12. Genotypes proportions

A Muller plot was used to visualize the emergence or disappearance of lineages specific to the origin of the initial virion suspension used to infect oysters. As previously described, the P0 viral inoculum was produced by pooling nine viral suspensions. To identify iSNVs specific to each origin, libraries from individuals sampled during the same mortality events within each basin were analyzed. Raw reads were filtered and trimmed as previously described, then aligned to the “artificial OsHV-1 major NR-consensus”. Variants were called using Freebayes and assigned unique identifiers as described above (Figure 1).

Because each sample originated from a specific location, each detected iSNV could be linked to its geographical origin. iSNVs identified throughout the EE of OsHV-1 µVar were grouped by origin and used to visualize their relative abundance using the MullerPlot package v0.13 (Farahpour et al., 2022) in R (Figure 1). Five categories were defined to track the dynamics of viral genomic backgrounds during experimental evolution: four categories corresponding to the distinct geographic origins composing the initial inoculum, and one additional “other” category grouping iSNVs that could not be confidently assigned to a single origin, likely reflecting shared polymorphisms among source populations.

## 3. Results

To investigate how OsHV-1 evolves during successive infection cycles, we conducted a multigenerational experimental evolution using *M. gigas* oysters. A single pool of virion suspension was used to infect specific-pathogen-free (SPF) oysters, which were subsequently collected from either oyster farming (FA) or non-farming areas (NFA) along the French Atlantic coast. This protocol was repeated over successive passages to simulate natural virion transmission under controlled environmental conditions.

In each passage, moribund oysters were sampled, and intra-host OsHV-1 µVar diversity was assessed through high-throughput sequencing. iSNVs were called to monitor changes in OsHV-1 µVar diversity and composition over time and across different host populations. Specifically, we examined patterns of iSNV accumulation, substitution biases, and lineage dynamics between oyster passages.

### 3.1. Evolution of survival rates and viral load in Pacific Oyster spat infected with OsHV-1 across passages

To assess the impact of OsHV-1 µVar infection on Pacific oyster spat, survival was monitored over 7 days post-infection (dpi) in both infected replicates and control tanks across three oyster populations. No mortality was observed in control tanks (*i.e.* ASW-injected oysters) throughout the experiment.

Cohabitation-based infection depends on effective within-host virion propagation. Challenges in maintaining reinfection across passages led to the termination of experimental evolution at the 13^th^ passage for the farming area (FA) population, the 18^th^ for the control (C) population, and the 28^th^ for the non-farming area (NFA) population.

Mortality rates varied across passages but showed an overall decreasing trend (Figure 3A and Figure S2A). At P0, mortality reached 88% in FA, 96% in C, and 84% in NFA oyster populations. In the C population, mortality decreased steadily to 42% at P15 before rising to 56% at P18, after which the experiment was stopped due to the absence of mortality at P19.

**Figure 3:**
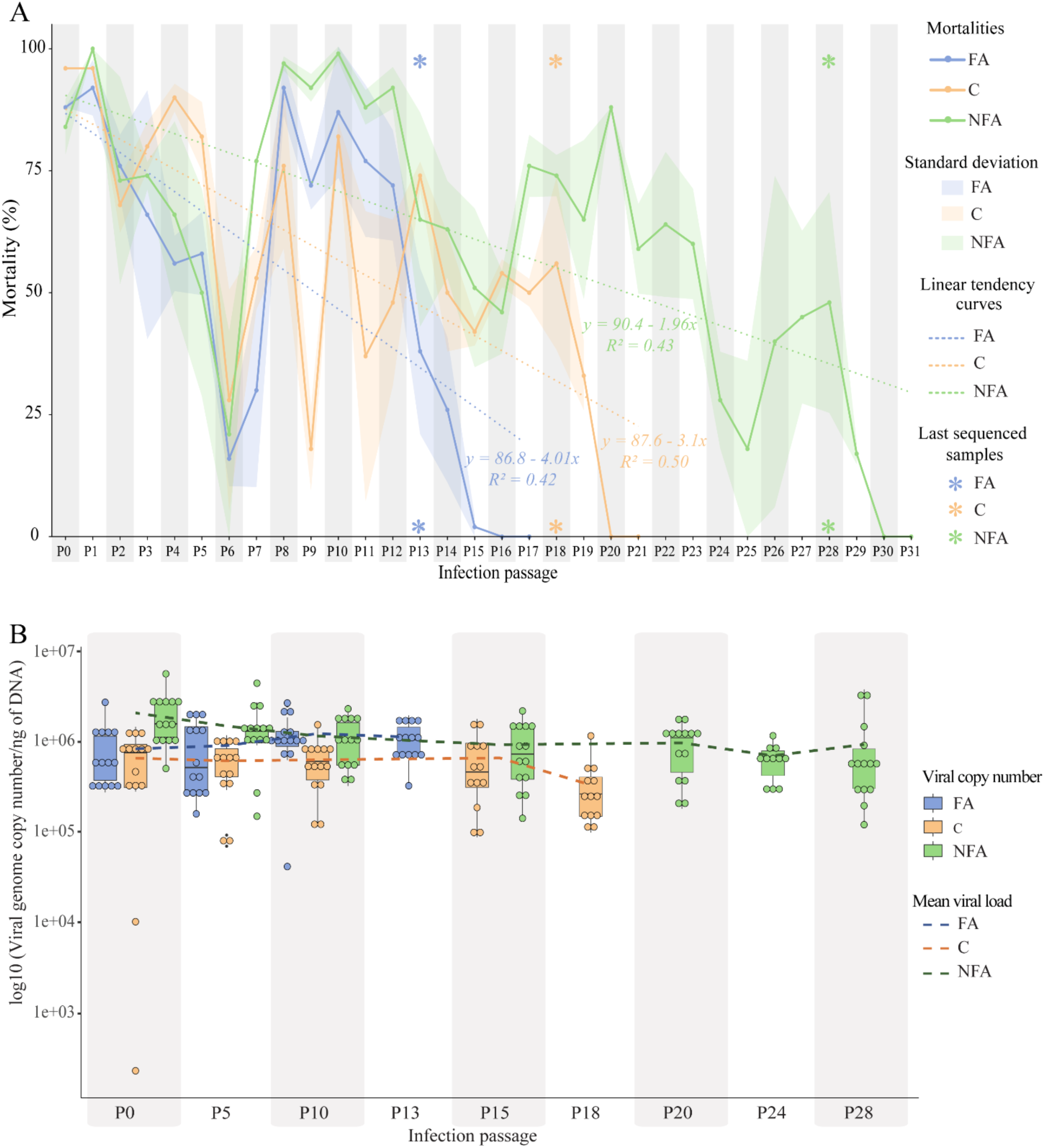
Oyster mortalities and virion load in the three oyster populations at passages 0, 5, 10, 13, 15, 18, 20, 24, and 28. A) Percentage of mortality observed at each passage in oysters from the Farming Area (FA, blue), Control (C, orange), and Non-Farming Area (NFA, green) populations. Mortality was monitored daily, and the proportion of dead oysters per passage is represented by solid lines. Dashed lines indicate the linear regression trend for each population across passages. Asterisks denote the final passage sequenced for each population. B) Individual viral genome copy numbers measured in infected oysters, expressed as log10-transformed viral genome copy number. Viral genome copy numbers were quantified from seven oysters per replicate per passage. Dots represent individual oysters, while box plots show the median, interquartile range, and variability within each group. Dotted lines connect mean viral genome copy numbers across passages for each population.

In contrast, FA and NFA populations exhibited more variable dynamics. In FA oysters, mortality declined to 58% at P5, then rose to 87% at P10, before falling sharply to 2%, which led to experimental termination. NFA oysters followed a similar pattern, with mortality initially decreasing to 50% at P5, peaking at 99% at P10, and subsequently fluctuating between 28% and 89%.

A global covariance analysis of mortality rates between replicates (Figure S2B) revealed a consistent correlation between the two replicates across populations, with correlation coefficients ranging from 0.70 to 0.83, supporting the robustness of the observed trends in survival over successive passages.

Viral genome copy number also varied among individuals and across oyster populations (Figure 3B). In FA oysters, the mean viral genome copy number increased slightly over passages, rising from 8.38×10^5^ ± 5.49×10^5^ viral genome copies/ng of DNA at P0 to 1.13×10^6^ ± 4.81×10^5^ viral genome copies/ng of DNA at P13 (Figure 3B). Conversely, C and NFA populations showed slight decreases in viral genome copy number across passages. In C oysters, viral genome copy number decreased from 6.62×10^5^ ± 4.49×10^5^ viral genome copies/ng of DNA at P0 to 3.34×10^5^ ± 2.54×10^5^ viral genome copies/ng of DNA at P18. For NFA oysters, viral genome copy number decreased from 2.09×10^6^ ± 1.20×10^6^ viral genome copies/ng of DNA at P0 to 9.35×10^5^ ± 1.11×10^6^ viral genome copies/ng of DNA at P28 (Figure 3B). Only selected passages are displayed to provide a clear overview of temporal trends, and when experimental evolution ended before P30 for a population, only the final available passage is shown.

ANOVA results revealed that infection passage had no significant effect on mortality (F(1,128) = 0.4054, p = 0.525464), while oyster population had a significant effect on mortality (F(1,128) = 0.12.5226, p = 0.000561) (Table 1). Similarly, viral genome copy number was significantly influenced by oyster population (F(1,215) = 1.0779e+10, p < 0.001), while infection passage had no significant effect (F(1, 3.0762e+08) = 0.7720, p = 0.37959). Together, these results indicate that mortality and viral genome copy number vary significantly among oyster populations and do not vary across infection passages.

**Table 1.** Summary of ANOVA results testing the effects of oyster population and infection passage on mortality and virion load. Mortality and viral genome copy number were analyzed as a function of oyster population and infection passage. Significant effects were considered if p < 0.05. DFn: degrees of freedom (numerator); DFd = degrees of freedom (denominator); F = F statistic.

|  |  | Sum of squares | DFn | DFd | F | P-value | Significance |
|---|---|---|---|---|---|---|---|
| Mortalities | Infection passages | 209.2 | 1 | 128 | 0.4054 | 0.525464 |  |
|  | Oyster population | 6463.9 | 1 | 128 | 12.5226 | 0.000561 | *** |
|  | Passages:Family | 1544.5 | 1 | 128 | 2.9921 | 0.086080 |  |
| Virion load | Infection passages | 4.3230e+11 | 1 | 3.0762e+08 | 0.7720 | 0.37959 |  |
|  | Oyster population | 1.0224e+13 | 1 | 1.0779e+10 | 18.2586 | 1.929e-05 | *** |
|  | Passages:Family | 2.6326e+12 | 1 | 3.1166e+08 | 4.7016 | 0.03013 | * |

### 3.2. Dynamics of substitutions and transition:transversion ratio

Studying substitution patterns across virion passages in EE is crucial to understand evolutionary dynamics, genetic diversity, and adaptation (Bromham, 2020). Based on the iSNVs identified from all sequencing libraries, we examined substitution dynamics across virion passages and oyster populations by analyzing ancestral iSNVs - defined as iSNVs already present in the initial inoculum - and *de novo* iSNVs - defined as iSNVs absent from the initial inoculum or observed in single individuals and arising during the experimental evolution process - counts and transition:transversion (ts:tv) ratios. A clear trend emerges across all three oyster populations, showing a rapid decline in the number of ancestral iSNVs in the OsHV-1 µVar genome over successive infection passages (Figure 4A). This decline is most pronounced in the FA population, with iSNVs counts dropping from 346 ± 26 iSNVs at P0 to 139 ± 59 iSNVs at P13 (Figure 4A). The number of ancestral iSNVs remains comparatively high in the NFA population (*e.g.* 329 ± 53 at P0 and 236 ± 12 at P28), with a slight increase observed at P10 (Figure 4A). *De novo* substitutions were systematically fewer than ancestral ones (*i.e.* typically fewer than 40 per passage across populations) and their numbers increased over successive infection passages (Figure 4C). For all three populations, most *de novo* iSNVs arise between P0 and P5, and remain relatively constant in subsequent passages.

**Figure 4.**
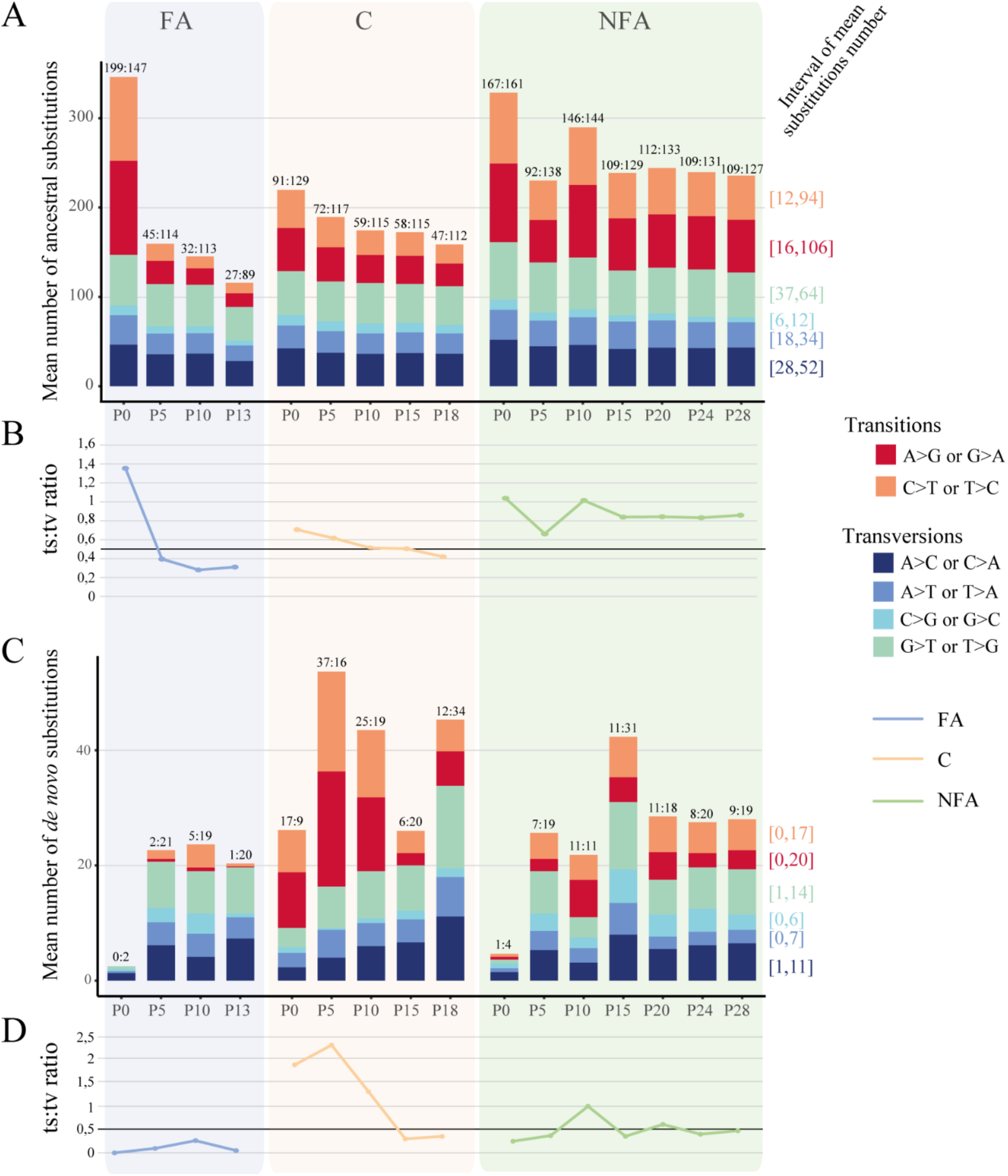
Substitution types and transition:transversion (ts:tv) ratio observed within OsHV-1 populations. A) Mean number of ancestral substitution types per passage and oyster population. Transitions (pyrimidine ↔ pyrimidine or purine ↔ purine) are shown in red to orange, and transversions (purine ↔ pyrimidine) in blue to green. Numbers above stacked bars indicate counts of transitions and transversions while numbers in brackets represent the mean per substitution type across samples. B) Variation of the ts:tv ratio across passages for each oyster population. Line colors correspond to the different oyster populations. C) Mean number of *de novo* substitution types per passage and oyster population. Colors and annotations follow the same scheme as in panel A. D) Variation of the ts:tv ratio for *de novo* substitutions across passages and populations.

A statistical analysis of the identified iSNVs was performed to assess the ts:tv ratio (Figure 4B and 4D). For ancestral iSNVs, the ts:tv ratio remained consistently between 1 and 0.4 across passages and populations, indicating a slight excess of transitions. The only exception was the FA population, where the ratio reached 1.26 at P0, reflecting a stronger transition bias, and then declined to below 0.4 from P5 onward, indicating a shift toward an excess of transversions (Figure 4B). In addition, the ts:tv ratio tended to decrease over passages in both the FA and C populations. For the NFA population, although the overall trend also points to a decline in the ts:tv ratio, a temporary increase was observed at P10. The ts:tv ratio for *de novo* iSNVs shows an almost opposite trend compared with ancestral iSNVs. In the FA population, values range from 0 to 0.4, whereas in the C population they span from 0.4 to 2.5, and from 0.4 to 1 in the NFA population (Figure 4D). Overall, the ratio tends to increase between P0 and P5 or P10, followed by a subsequent decline.

As observed with ts:tv ratio analysis, intra-individual nucleotide diversity (π) varied markedly across passages and among viral families (Figure S3). At passage P0, π was highest in FA and NFA, whereas C displayed lower initial diversity. A sharp decline in diversity was observed at P5 for all families, converging toward similar π values (∼0.025). At P10, contrasting dynamics emerged: NFA exhibited a pronounced and transient increase in nucleotide diversity, reaching the highest value observed across the experiment, while FA and C remained comparatively stable with moderate π levels. From P15 onward, diversity decreased and stabilized in all families, converging toward similar low values by the final passages (P25–P30). The shaded ribbons indicate greater variability among replicates at early passages, particularly for FA at P0 and NFA at P10, followed by reduced dispersion at later passages. Overall, the figure highlights distinct early diversification dynamics among families, followed by progressive homogenization of intra-individual genetic diversity over successive passages.

### 3.3. Unveiling OsHV-1 population structure and anomalies in experimental evolution through iSNVs data analysis and principal component projection

To investigate OsHV-1 µVar population structure across passages and oyster populations, we performed a Principal Component Analysis (PCA) on iSNVs frequency data, incorporating metadata on host population, infection passage, and replicates (Figure 5). To address potential concerns about restricting the analysis to only two components, we further extended the PCA to the first 25 components (Figure S1). This extended analysis confirms that the major clustering patterns observed in Figure 5 are consistent with the broader genomic variation, while also showing that subsequent components contribute relatively little to the explained variance. We therefore retained the first two components in the main figure for clarity, as they best capture the main biological signals of infection progression and host population differences. The first principal component accounted for 12.61% of the variance, while the second component explained 9.89% of the variance (Figure 5). The first component reflects infection progression over passages, whereas the second is associated with oyster population or susceptibility traits. The plot reveals three distinct clusters corresponding to the interplay between oyster populations and infection passages.

**Figure 5.**
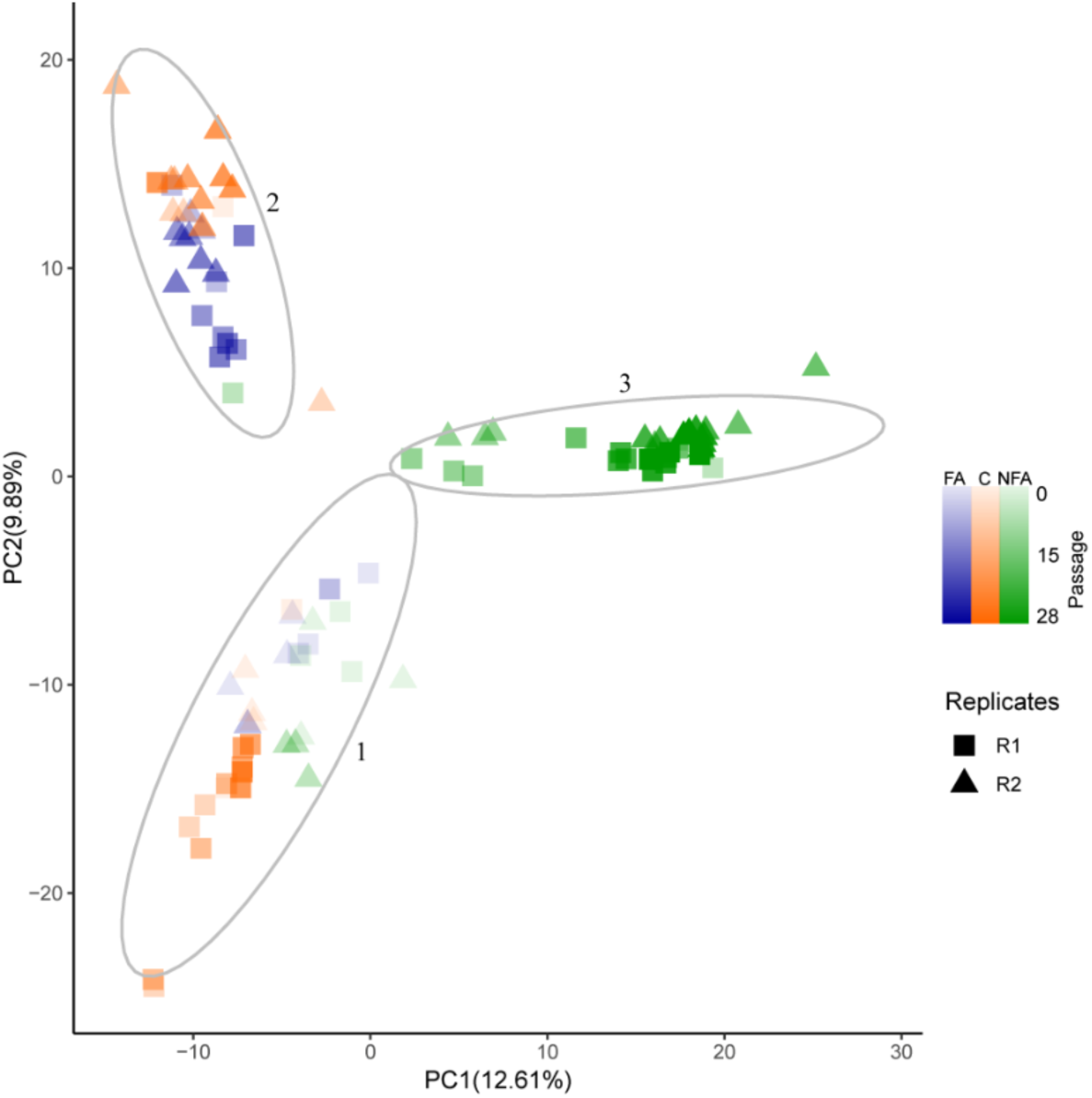
Principal Component Analysis (PCA) of OsHV-1 genetic variation among viral populations from different oyster populations and infection passages. PCA was performed on iSNVs detected within OsHV-1 genomes to investigate patterns of genetic differentiation among viral populations and infection passages. Each point represents an individual viral genome, projected onto the first two principal components (PC1and PC2). Colors indicate the host oyster population: blue for FA (Farming Area), orange for C (Control), and green for NFA (Non-Farming Area). Replicates are distinguished by point shapes (squares for replicate 1 and triangles for replicate 2), and infection passages are represented by a color gradient. The ellipses delineate three major clusters of genetically related OsHV-1 µVar populations, labeled Clusters 1 to 3.

The first cluster includes samples from all oyster populations at P0, along with those from replicate 1 of the C population at P0, P5, P10, P15, and P18 (Figure 5, group 1) suggesting that the initial OsHV-1 µVar diversity was maintained across passages in replicate 1 of the C population.

The second cluster (Figure 5, group 2) includes samples mainly from the FA population at P5, P10, and P13, as well as from replicate 2 of the C population at P5, P10, P15, and P18.

Finally, the third cluster comprises samples from the NFA population, which includes samples from P15 and P28 (Figure 5).

### 3.4. Detection of regions under selection pressure and allelic frequency trajectories

To detect genomic regions under selection, we applied the Cochran-Mantel-Haenszel (CMH) test independently to iSNVs identified in each oyster population, focusing on allele frequency changes between the initial and evolved passages. The CMH test is widely used to detect selection pressures by analyzing allele frequency shifts in experimental evolution and time-series data (Spitzer et al., 2020).

The CMH test, applied to iSNVs identified in each OsHV-1 µVar population from the three oyster populations independently (Figure 6), showed that no genomic regions were significantly affected by selection in the FA (Figure 6A) and C (Figure 6B) populations. In contrast, the NFA population exhibited 117 candidate iSNVs under selection (Figure 6C).

**Figure 6:**
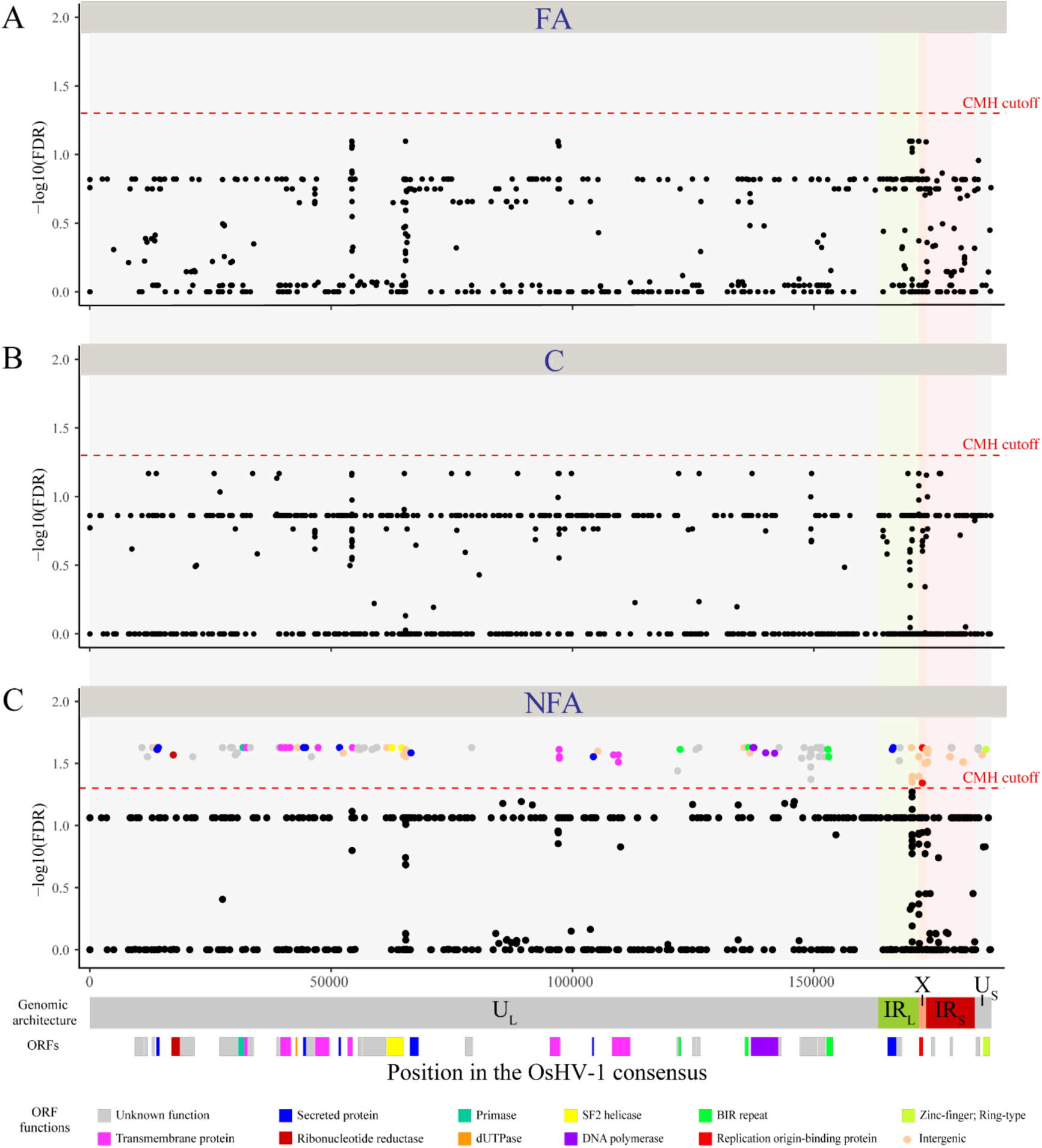
Genomic distribution of candidate iSNVs under selection in oyster populations. The scatter plots show the negative log10-transformed p-values of iSNVs along the OsHV-1 genome for A) Farming Area, B) Control and C) Non-Farming Area experimental oyster populations. Each point represents an iSNV identified by comparing OsHV-1 µVar populations from the founder and evolved passages using the Cochran–Mantel–Haenszel (CMH) test. The red dashed line indicates the significance threshold corresponding to a 5% false discovery rate (CMH cutoff). iSNVs above this threshold are considered candidates for selection and are color-coded based on their potential functional impact. The OsHV-1 genome structure is shown below the plots, including unique regions (U_L_ and U_S_) and inverted repeat regions (IR_L_ and IR_S_). Colored blocks correspond to Open Reading Frames (ORFs) containing candidate iSNVs, with colors reflecting the predicted function or protein domain of each ORF.

Among the 117 candidate iSNVs identified in the NFA population, 73 corresponded to ancestral variants already present in the inoculum, whereas 44 were de novo mutations that emerged exclusively in the NFA population (Table 2). Ancestral iSNVs were predominantly transitions (60 transitions among 73 iSNVs), with only 7 transversions, 5 deletions, and 1 insertion. In contrast, *de novo* iSNVs were mainly transversions (33 transversions among 44 *de novo* iSNVs), along with 5 transitions, 3 deletions, and 3 insertions. These results indicate that most selected variants in the NFA population originated from standing genetic variation, while a substantial fraction arose during experimental evolution.

**Table 2.** Distribution of candidate iSNVs under selection in the NFA population according to their status and mutation type. The table summarizes the 117 iSNVs identified by the CMH test in the NFA population, classified as ancestral (present in the inoculum) or *de novo* (emerged during experimental evolution process), and further categorized by mutation type (transition, transversion, deletion, insertion). Totals are provided for each category.

|  | Transition | Transversion | Deletion | Insertion | **Total** |
|---|---|---|---|---|---|
| Ancestral iSNVs | 60 | 7 | 5 | 1 | **73** |
| *De novo* iSNVs | 5 | 33 | 3 | 3 | **44** |
| **Total** | **65** | **40** | **8** | **4** | **117** |

These iSNVs were distributed across the OsHV-1 genome, with a concentration in repeated regions and in the U_L_ region between 50 kb and 80 kb. More precisely, 28 potentially selected positions were located in intergenic regions, including one in the stem loop, and 89 iSNVs were located within ORFs (Figure 6C, Table S3).

Among the 89 iSNVs located within ORFs, 58 were found in ORFs of unknown function, and 12 in ORFs encoding transmembrane proteins (Table S3). In addition, 8 iSNVs were located in ORFs encoding secreted proteins, four in ORF 100 coding for a DNA polymerase, three within ORFs encoding Zinc-finger, Ring-type proteins, two within ORFs encoding BIR repeat motifs, one within ORF115 associated with an Origin-binding protein, and the final iSNV within ORF20 encoding a ribonucleotide reductase (Table S3).

An allele frequency trajectory (AFT) plot was used to track changes in the frequencies of candidate iSNVs identified in the NFA oyster population, providing insights into evolutionary dynamics and signals of positive selection (Barghi et al., 2020). The analysis was conducted separately for both replicates (Figure 7).

**Figure 7:**
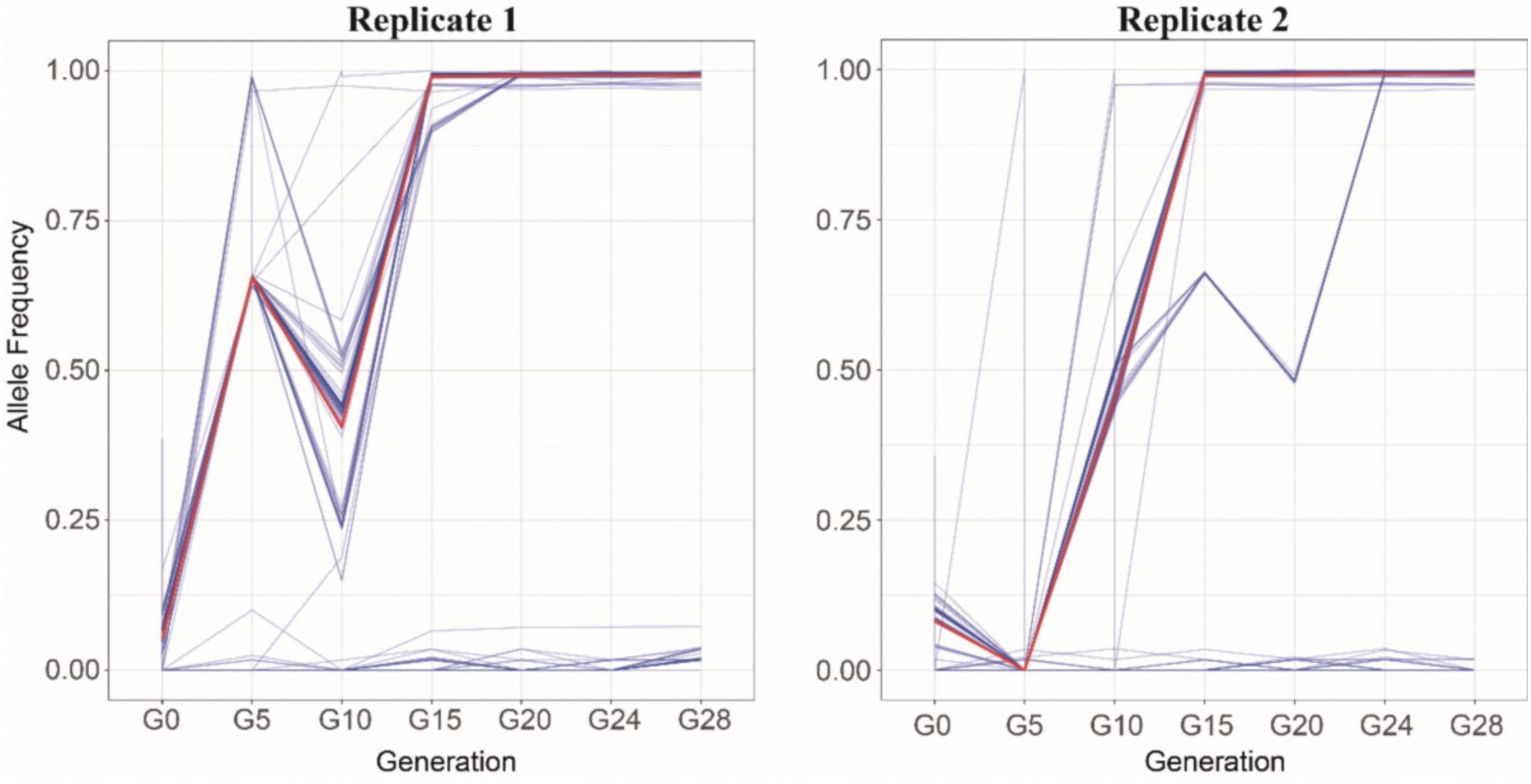
Temporal trajectories of allelic frequencies for top 100 candidate OsHV-1 iSNVs during serial infections in the NFA oyster population. Each blue line represents the allelic frequency trajectory of a specific iSNV across passages. Red line represents the median allelic frequency trajectory.

The AFT plot shows a general trend of increasing allele frequencies, often leading to fixation of selected alleles in both replicates. In replicate 1, allele frequencies of candidate iSNVs rise from P0 to P5, drop between P5 and P10, and finally increase again between P10 and P15 until fixation (Figure 7A). A similar trend is observed in replicate 2, but only for 77 iSNVs (Figure 7B). Moreover, 66 iSNVs in replicate 1 and 26 iSNVs in replicate 2, shown at the bottom of the plot, also exhibit increasing frequencies, though to a lesser extent (Figure 7A and B).

To further investigate potential linkage among candidate iSNVs, we identified 74 local haplotype blocks, defined as groups of SNPs separated by less than 300 nucleotides (Table S4). Most blocks contained a single iSNV, whereas several blocks harbored multiple co-occurring variants. The largest block, Block_57 (positions 149127–149494), contained up to nine iSNVs. These multi-iSNV blocks reveal genomic regions where mutations consistently co-occurred and exhibited coordinated allele frequency changes across replicates.

### 3.5. Relative abundance of iSNVs from specific origin

To monitor the persistence and dynamics of viral lineages from specific origins across passages of infection, a Muller plot was generated (Figure 8). The results show that multiple lineages from different origins coexist from the start to the end of the experiment within each oyster population. At passage 0, lineages specific to MO and LEU had a higher relative abundance (RA) than those from ARC and BR. This general trend was maintained across passages of infections in all oyster populations, though RA varied depending on the population. In the FA population, OsHV-1 iSNVs specific to MO are present at a RA of 0.2 at P0, which increased to 0.4 by P10 then slightly declined. A similar trend was observed for iSNVs from ARC and BR, but with much lower RA values (between 0.02 and 2.0). In contrast, the RA of LEU-specific iSNVs declined from 0.4 at P0 to 0.1 at P5.

**Figure 8.**
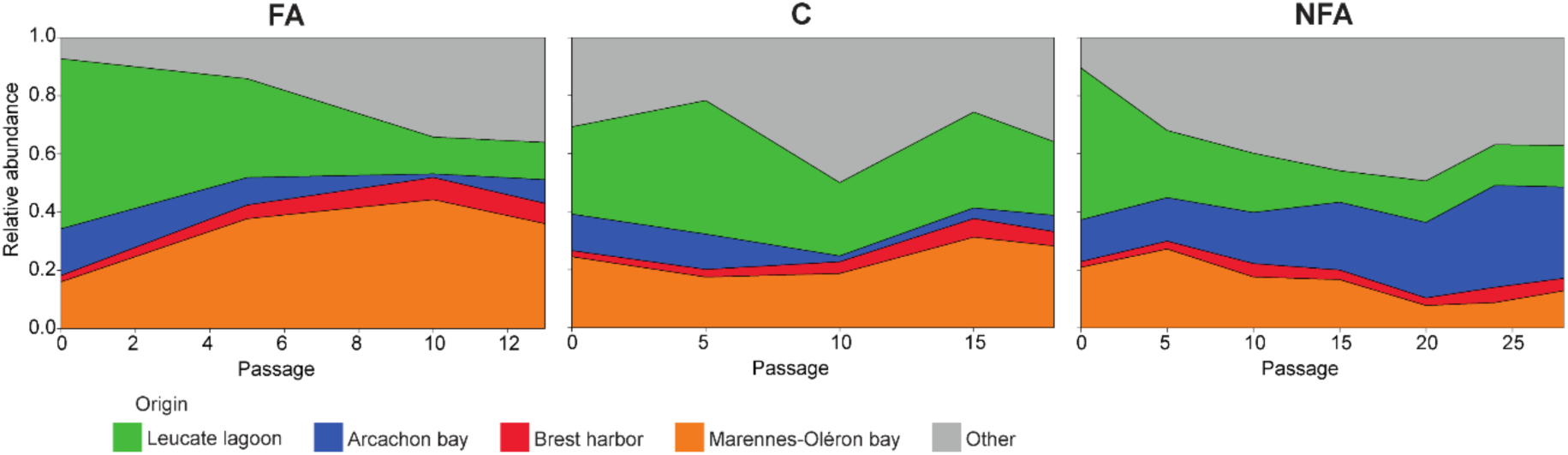
Muller plot representing the relative abundance of viral iSNVs collected from specific origins for each oyster population during the experimental evolution. The relative abundance of viral SNPs is color-coded according to their geographical origin of the viral suspension: the orange area corresponds to lineages collected specifically in MO, the red area corresponds to lineages collected specifically in BR, the blue area corresponds to lineages collected specifically in ARC, the green area corresponds to lineages collected specifically in LEU, and the grey area corresponds to lineages that appeared during the experimental evolution.

New iSNVs, absent from all initial viral suspensions, began to emerge during the experiment. At P0, their RA was low in both farming and non-farming populations (RA = 0.1), but increased over passages. In contrast, in the control population, the RA of these “other” iSNVs started at 0.3, dropped to 0.2 by P5, rose to 0.5 at P10, then declined again to 0.2 by P15.

## 4. Discussion

To mitigate the impact of infectious diseases, it is essential to understand the mechanisms shaping the evolution and spread of pathogenic microorganisms (Gu et al., 2021; NIH, 2007). Variation in host resistance plays a major role in driving these processes (Gandon & Michalakis, 2000; Kubinak & Potts, 2013). This study provides valuable insights into the evolutionary dynamics of OsHV-1 and its interactions with Pacific oysters, focusing on how the virus adapts and diversifies in response to host genetic backgrounds. These findings are crucial to address the devastating impact of OsHV-1 on oyster farming. The similar intercepts observed in the mortality regression models indicate that early infection passages (P0 to P2) resulted in comparable mortality levels across experimental conditions. This pattern is consistent with the use of a heterogeneous mixture of viral inocula from multiple geographic origins, in contrast to previous assessments of resistance based on a single-origin inoculum from the Marennes– Oléron basin. Such viral mixtures likely include genetically and phenotypically diverse variants, generating a strong acute infection that can mask subtle differences in host resistance at the onset of infection. Accordingly, similar early mortality values likely reflect a shared initial susceptibility to a complex and highly virulent viral challenge rather than equivalent resistance levels. In contrast, resistance- or tolerance-associated differences emerge through the temporal dynamics of mortality across passages, as indicated by the divergence in regression slopes, emphasizing that resistance should be interpreted as a dynamic trait expressed over time.

During this experimental evolution, we observed a decrease in both oyster mortalities and viral genome copy numbers across successive passages of OsHV-1 infection. Interestingly, despite the distinct infection histories of the three oyster populations, mortality patterns were unexpectedly similar during the early passages of infection, contrary to the anticipated gradient. This convergence in mortality rates suggests that the initial infection dynamics may have been strongly influenced by experimental conditions rather than by prior host exposure history. For instance, standardized infection doses and cohabitation settings could have minimized differences in host susceptibility or viral particle virulence between populations. The decline in mortalities and viral genome copy number results from several, non-exclusive factors. First, it can be explained by the selection of less virulent viral lineages. Host-pathogen interactions are shaped by a balance between virulence and transmission (Anderson & May, 1982). In conditions with limited host numbers, such as in our setup with 25 oysters per tank, rapid host death could threaten virion persistence. A plausible strategy for OsHV-1 may be to reduce virulence, and maintaining a stable host population while ensuring replication. This trade-off between virulence and transmission is well-documented, with higher transmission usually associated with lower virulence, and conversely, excessive virulence reduces transmission (Anderson & May, 1982; Gandon & Michalakis, 2000; Kun et al., 2023). Comparable results were reported in EE study involving tobacco etch potyvirus (TEV) in plants, where viral genome copy number and virulence decreased across passages, strongly influenced by host lineage (Cuevas et al., 2015). However, virulence does not necessarily decrease during host–virus co-evolution. In some systems, virulence has increased over time, as observed in Marek’s disease virus (Trimpert et al., 2017, 2019), illustrating that the direction of virulence evolution is context-dependent.

Secondly, the reduction in mortality rates can be attributed to a lack of complementation between lineages (Froissart et al., 2004; Montville et al., 2005). Host infection rarely occurs with a single lineage, but often involves co-infection, with multiple viral lineages contributing to the infection and associated symptoms. Certain lineages carrying disadvantageous mutations can be complemented by co-infecting high-fitness lineages, allowing low-fitness lineages to gain a phenotypic advantage (Froissart et al., 2004; Montville et al., 2005). In our study, where genetic diversity has likely been drastically reduced, possibly due to a bottleneck effect, some lineages may have been purged, limiting complementation and reducing virulence over successive passages. Bottlenecks can drastically reduce genetic diversity by eliminating rare lineages and fixing certain alleles through genetic drift (Zwart & Elena, 2015). In viral populations, this can compromise cooperative interactions between lineages, diminishing overall viral fitness and reduction in mortality rates (Zwart & Elena, 2015).

A third hypothesis to explain declines in mortality rates and viral genome copy numbers is a temporal shift in the peak of viral particle release into seawater across infection passages. OsHV-1 excretion peaks between 24 and 48 hours post-infection (hpi) (Delmotte et al., 2020), but timing can vary with oyster susceptibility, initial viral genome copy number, donor numbers and age, food availability, water temperature, and salinity (Dégremont, 2011; Delmotte et al., 2020; Evans et al., 2017; Pernet et al., 2018, 2018; Petton et al., 2015; Schikorski et al., 2011). Consequently, the viral genome copy number would be high when cohabitation begins and progressively decreasing over time. In our experiment, successive infections involved oysters from the same cohort, whose age increased across passages, potentially altering susceptibility and timing of viral excretion peak. Cohabitation infections can also lead to fewer fatalities due to progressive decline in viral particles number across cycles (Cain et al., 2021). While it may not be the optimal method for EE, successive infection by cohabitation was chosen to mimic the natural infection of oysters by OsHV-1 in the field.

A fourth complementary hypothesis relates to latent or persistent OsHV-1 infections. OsHV-1 may establish a latent or persistent state in oysters, allowing persistence without acute disease (Divilov et al., 2024), as seen in other herpesviruses (Dotto-Maurel et al., 2025; White et al., 2012). FA oysters indeed exhibited high viral genome copy numbers even during low mortality periods, suggesting a shift toward non-lytic or less virulent states, potentially mediated by host immune regulation or viral adaptation favoring long-term coexistence (Divilov et al., 2024). In contrast, the C population show lower viral genome copy numbers than FA population indicating that viral replication efficiency, rather than immediate host mortality, can vary among populations. These patterns may reflect differences in host immune responses or in the balance between lytic and persistent infection. Although our experimental design did not include latency-specific assays, these observations highlight the need to further investigate the latent potential of OsHV-1 under recurrent exposures and genetically selected oyster populations.

Genetic mutations, including transitions and transversions, shape viral evolution by contributing to genetic diversity and adaptation and molecular evolutionary hypothesis suggests that natural selection promotes amino acid substitutions mostly through transitions (Stoltzfus & Norris, 2016). Importantly, although the three experimental populations were infected with an identical viral inoculum, substantial differences in standing genetic diversity were already observed at P0. This apparent discrepancy likely reflects strong stochastic effects during the earliest stages of infection, including transmission bottlenecks and host-specific sampling of the viral mutant spectrum. Because each oyster constitutes an independent biological replicate, only a subset of the virions present in the inoculum may successfully initiate replication within a given host. Consequently, early interindividual variation in iSNV composition can generate marked differences in apparent standing diversity among populations at P0, even in the absence of differences in the source inoculum. These early founder effects likely shaped subsequent evolutionary trajectories across passages. The decline in intra-individual single nucleotide variants (iSNVs) and in the ancestral transition:transversion (ts:tv) ratio across passages suggest progressive genetic homogenization. Nevertheless, *de novo* iSNVs increased sharply at P5 before stabilizing, indicating ongoing mutation emergence despite reduced overall diversity. Interpretation must consider potential technical artefacts, including sequencing errors or library preparation bias (e.g. oxidative DNA damage generate G|T and C|A transversions), low frequency false-positive calls particularly for viral quasispecies, or misalignments in repetitive, low complexity of poor mapability genomic regions can produce spurious iSNVs, often enriched in transversions (Jensen et al., 2021; Lenz et al., 2014; López-Muñoz et al., 2021; Phan et al., 2015; Poetsch, 2020; Van Poelvoorde et al., 2022). Biologically, the fidelity of the OsHV-1 DNA polymerase, as well as host-mediated processes such as oxidative stress or deamination, could also favor transversions (Mourier et al., 2021). Natural selection may further modulate which lineages persist. Analyses based on a global consensus genome (i.e. 126 viral NR-genomes) may additionally introduce reference biases affecting iSNVs detection. Moreover, because these reconstructed genomes were generated from short-read sequencing data (2 × 150 bp), they should be interpreted as sample-specific consensus assemblies rather than fully phased 200 kb viral haplotypes, as short reads inherently limit the resolution of long-range linkage among distant variants within mixed viral populations. Indeed, the ts:tv ratios observed here are lower than those reported for humans (∼2) or for Varicella-Zoster virus (∼2.3), consistent with technical influences, but they may also reflect virus- or host-specific mutation biases (Jeon et al., 2016; Wang et al., 2015). These observations align with patterns in other viral systems, where bottlenecks and genetic drift or selective sweeps reduce diversity across passages (Kutnjak et al., 2017; Li & Roossinck, 2004). Given the context of OsHV-1 infecting oyster of distinct genetic backgrounds, a combination of technical artefacts, host-mediated processes, and genuine evolutionary bottlenecks likely contributed to reduced viral lineages.

PCA analysis of iSNV frequencies revealed clusters corresponding to oyster populations and infection passages, indicating genetic differentiation and adaptability. Notably, clustering of NFA and FA individuals was clearly delineated, while control oysters exhibited intermediate clustering, suggesting susceptibility-dependent evolution (Kubinak & Potts, 2013). Within the NFA oyster population, distinct clusters emerged, indicating evolutionary changes across infection passages in response to selective pressures. The relative clustering homogeneity observed for NFA samples in the PCA is consistent with the substitution dynamics, where ancestral iSNVs remained abundant across passages while *de novo* substitutions accumulated only moderately. This suggests that NFA viral populations retained a large proportion of the initial standing diversity and evolved along relatively similar trajectories across replicates. In contrast, the separation between the two C replicates in the PCA likely reflects replicate-specific changes in iSNV composition, including differential retention of ancestral variants and distinct patterns of *de novo* substitution accumulation across passages.

The dynamics of iSNVs revealed clear differences among viral lineages of distinct geographic origins, highlighting how host environment and exposure history shape OsHV-1 diversity and persistence. Throughout the experiment, iSNVs from MO remained dominant in all oyster populations, particularly in those from FA and in control oysters, whereas iSNVs originating from ARC progressively increased in relative abundance within NFA oysters. This contrasting pattern likely reflects both the epidemiological history and ecological context of the host populations. All viral lineages used in the experiment originated from farming areas and were initially sampled during mortality events, suggesting that NFA oysters represent the most geographically and epidemiologically distant hosts from these viral sources. The expansion of ARC-like iSNVs in NFA oysters could indicate that these hosts, having experienced limited or no prior exposure to OsHV-1 lineages from ARC, and impose weaker selective constraints on viral replication. Such relaxed selection may allow for the persistence or even amplification of lineages that are not favored in aquaculture environments, where repeated exposure and strong host-virus coevolution tend to filter viral diversity (Kennedy et al., 2016). Conversely, the predominance of MO-like iSNVs in FA oysters likely results from repeated selective pressures imposed by ongoing viral circulation in dense farming contexts, favoring lineages already adapted to these hosts (Kennedy et al., 2016).

Interestingly, new iSNVs that were absent from all initial inocula also emerged during the course of the experiment, suggesting ongoing within-host diversification and potential *de novo* mutation events. The accumulation of these new variants, particularly in NFA oysters, may reflect broader environmental tolerance or increased mutational persistence under relaxed selective pressure. In contrast, the lower diversity observed in FA oysters supports the idea that repeated exposure to similar viral lineages and stronger immune selection in farmed populations drive the convergence toward dominant MO-like lineages.

The identification of 117 candidate selection signatures in the OsHV-1 genomes and allelic frequency trajectories (AFT) plots in the NFA population, provides strong evidence of selective pressures during experimental evolution. Allele frequency increases and fixation indicate positive selection (Barghi et al., 2020). Selection was detected exclusively in NFA oysters, suggesting enhanced viral replication that may increase mutation rates and facilitate the fixation of advantageous alleles in these more susceptible hosts (Dégremont, 2011; Elena & Sanjuán, 2005; Peck & Lauring, 2018). Conversely, no selection signals were detected in FA oysters, despite their reduced OsHV-1 diversity, supporting the influence of bottleneck and drift effects. Given that viral evolution are typically shaped by environmental changes and host immune pressures acting as selective filters to constrain viral diversity (Kubinak & Potts, 2013), this absence of detectable selection in FA oysters is unexpected, suggesting that stochastic processes such as bottlenecks and genetic drift dominated viral evolution in these populations. This may result from the repeated transmission and amplification of a limited number of viral lineages within farming environments, reducing opportunities for adaptive lineages to emerge and spread.

Selection signatures were distributed across the viral genome, particularly within repetitive regions and in the U_L_ region. Although the use of short-read sequencing prevented reconstruction of complete long-range haplotypes, the identification of local haplotype blocks provides a first approximation of the underlying haplotype structure within the viral population. The recurrent co-occurrence of multiple iSNVs within specific genomic regions suggests linked evolutionary dynamics and may reflect selection-driven linkage or the persistence of partially associated viral genetic backgrounds, particularly in the NFA population. Such a widespread pattern likely reflects a combination of immune-driven and relaxed selection processes acting in under contrasting host environments. According to the “relaxed selection hypothesis” (Mikonranta et al., 2012), a neutral virulence-related trait in environments where host density and immune pressure are low, lead to alteration or loss through the random accumulation of mutations rather than active negative selection. In the context of OsHV-1, NFA oysters, characterized by reduced host density and weaker immune activation, may thus favor the persistence of neutral or mildly deleterious mutations, potentially giving rise to attenuated viral lineages. This relaxation of selective constraints could therefore contribute to the progressive decline in mortality rates observed during the experiment, acting in parallel with adaptive processes occurring in farming environments.

These selection signals were further enriched in ORFs encoding proteins critical for viral attachment, replication, packaging, and membrane functions. Viral entry proteins, even with single amino acid changes, can alter infection success across hosts with different susceptibilities (van Sluijs et al., 2017). Acquired, duplicated or lost domains in Herpesviridae evolution, including envelope (such as membrane glycoproteins and transmembrane receptors), auxiliary domains (like Zinc-finger, RING type, dinucleoside kinase), and modulation domains (such as Interleukin, Interferon-regulatory factor, or Zinc-finger), enhance specific binding and immune evasion (Brito & Pinney, 2020). In OsHV-1, most selection signals were within these domains, indicating adaptation of essential proteins to host genetic backgrounds. Two previous studies support immune-driven adaptation in OsHV-1 across host species and environments (Delmotte et al., 2020; Pelletier et al., 2025). Based on the results obtained in this study, it is evident that adaptation can also occur at a finer scale within the host’s genetic background, which is influenced in this case by the geographic origin of the parental individuals (*i.e.* farmed or not farmed area).

## 5. Conclusion

This study provides valuable insights into the evolutionary driving mechanisms of OsHV-1 and its interactions with Pacific oysters, with a particular emphasis on viral adaptation and diversification in response to host infection susceptibility. Employing an experimental evolution approach, it sheds light on the dynamic nature of host-virus interactions, the potential for viral adaptation, and the possible role of genetic diversity in shaping the outcome of these interactions. The nucleotide evolution of OsHV-1, particularly a decrease in transition and transversion numbers, appears to be mainly driven by both genetic drift and positive selection linked to the oyster genetic background, resulting in a reduction in OsHV-1 diversity and the fixation of specific alleles within ORFs dedicated to host-virus interactions and virion functional maintenance. Further research into the functional consequences of these changes is necessary. These findings contribute to our understanding of the mechanisms governing viral evolution and hold significant implications for mitigating the devastating impacts of OsHV-1 on oyster farming industries. Future studies should continue to explore the intricate dynamics of host-virus interactions and their implications for viral evolution and disease management.

## Data availability

Raw data have been deposited on the SRA database under Bioproject PRJNA1216400 accession numbers SAMN46433918 to SAMN46434013 for future reference and accessibility (Table S2). All the scripts are available <u>at</u> https://gitlab.ifremer.fr/asim/experimental_evolution_public, at https://github.com/cpelleti/experimental_evolution_OsHV-1 and in the Zenodo platform (DOI: 10.5281/zenodo.24394048 ; https://zenodo.org/records/24394048)

## Acknowledgment

We thank the staff of the Ifremer station at La Tremblade (ASIM), Frédéric Girardin and his team (Plateforme des Mollusques Marins de La Tremblade PMMLT), and Virginie François and his team (Plateforme des Mollusques Marins de Bouin PMMB). We also thank the SEBIMER team for maintaining bioinformatics tools.

C.P. was financially supported by a grant from the Ifremer Scientific Board and the Nouvelle-Aquitaine region. The present study was supported by scientific direction of Ifremer in context of the GT Huître project and by the FEAMP Gestinnov (FFEA470020FA1000007) GESTINNOV project and by DGAL (French General Directorate for Food) through the National Reference Laboratory for Mollusc Diseases and by the European-Union Reference Laboratory for Mollusc Diseases, Ifremer, La Tremblade. We sincerely thank the French National Research Agency (ANR) for the time and support dedicated to the development of this work (ANR-23-CE35-0009). The authors acknowledge the Pôle de Calcul et de Données Marines (PCDM; https://wwz.ifremer.fr/en/Research-Technology/Research-Infrastructures/Digital-infrastructures/Computation-Centre) for providing DATARMOR computing and storage resources. A preprint version of this article has been peer-reviewed and recommended by PCI Evol Biol (https://doi.org/10.24072/pci.evolbiol.100858). The funders had no role in study design, data collection and interpretation, or the decision to submit the work for publication.

C.P. and co-authors thank J.M.E. for the discussions and exchanges. We would like to thank Bruno Petton and his team PHYTNESS for sampling viral particles in the Brittany area, as well as Johan Vieira and the CAPENA team for sampling OsHV-1 in Arcachon Bay. B.M., N.F., and C.P. designed the study; L.D., B.M., and J.V.D. provided the biological materials; N.F., M.H., M.M., and C.P. collected and processed the samples. C.P. performed the bioinformatics analysis of the Illumina data. B.M., G.C., M.J., J.V.D., L.D., and C.P. drafted the manuscript. All authors read and approved the final version of the manuscript.

We declare that we have no competing interests.

