## Supplementary figures and images for "Experimentally mimicking long term infections of *Magallana gigas* with the OsHV-1 virus reveals evolution through positive selection"

### Supplemental Figure 1

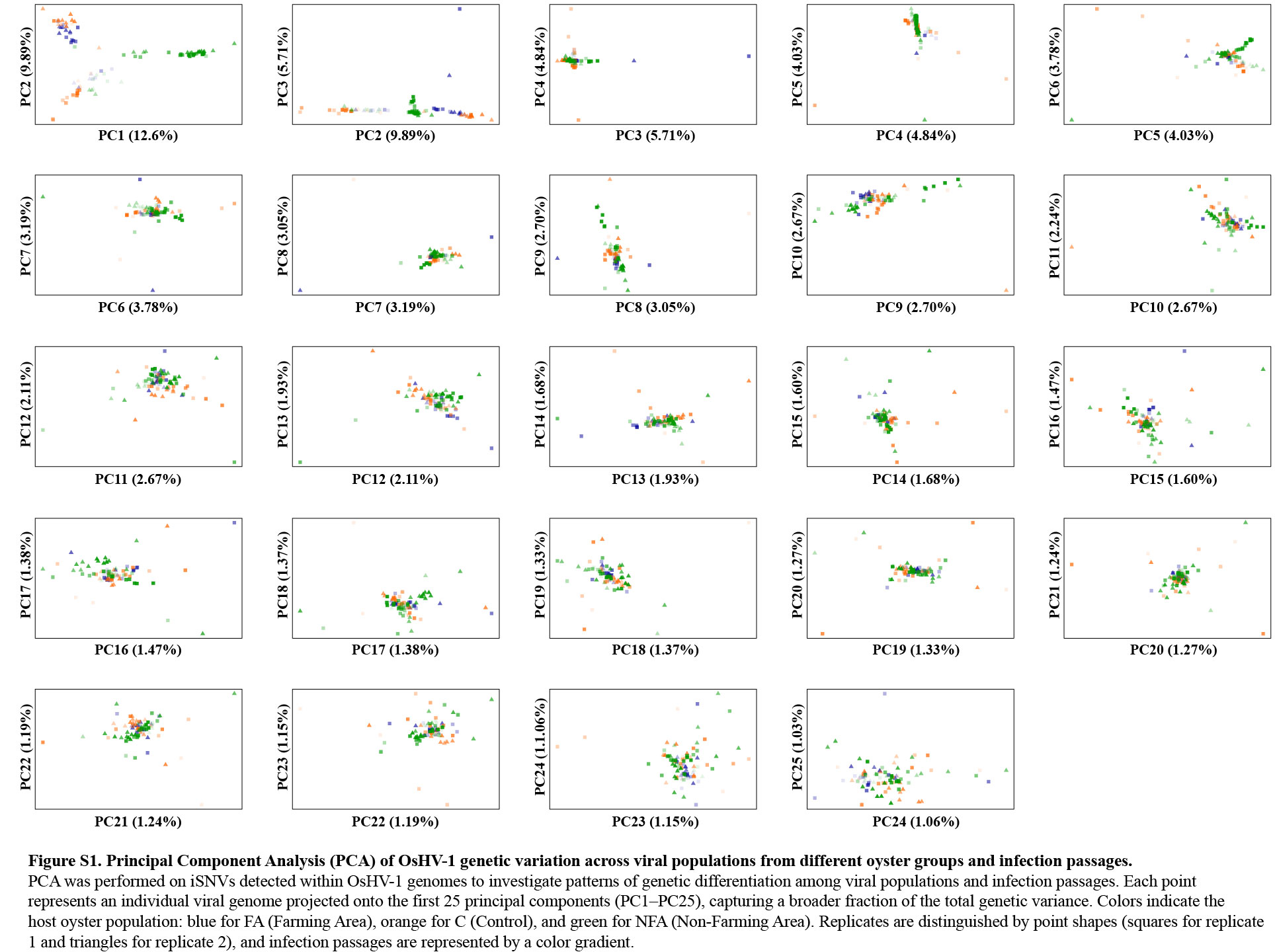

### Supplemental Figure 2

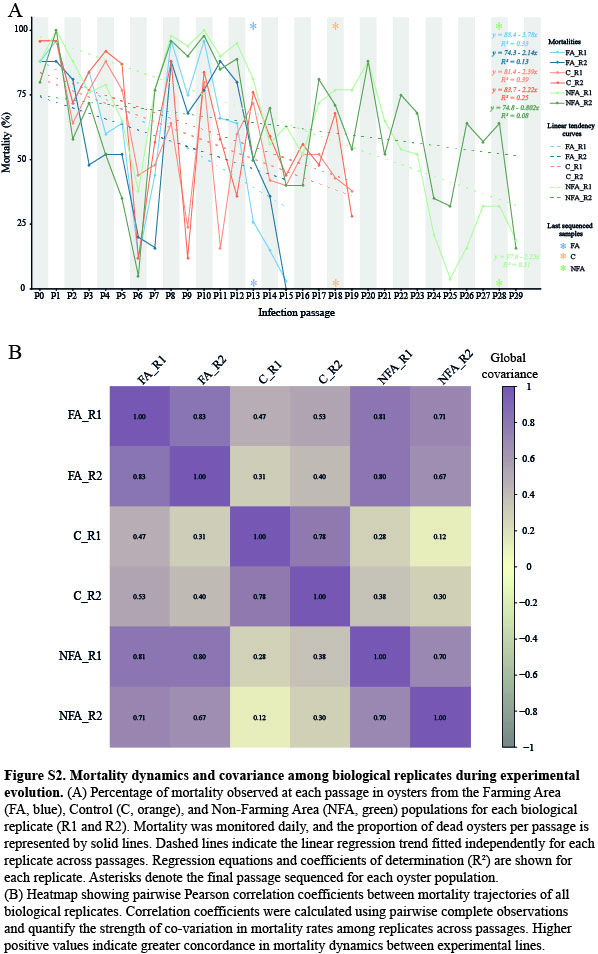

### Supplemental Figure 3

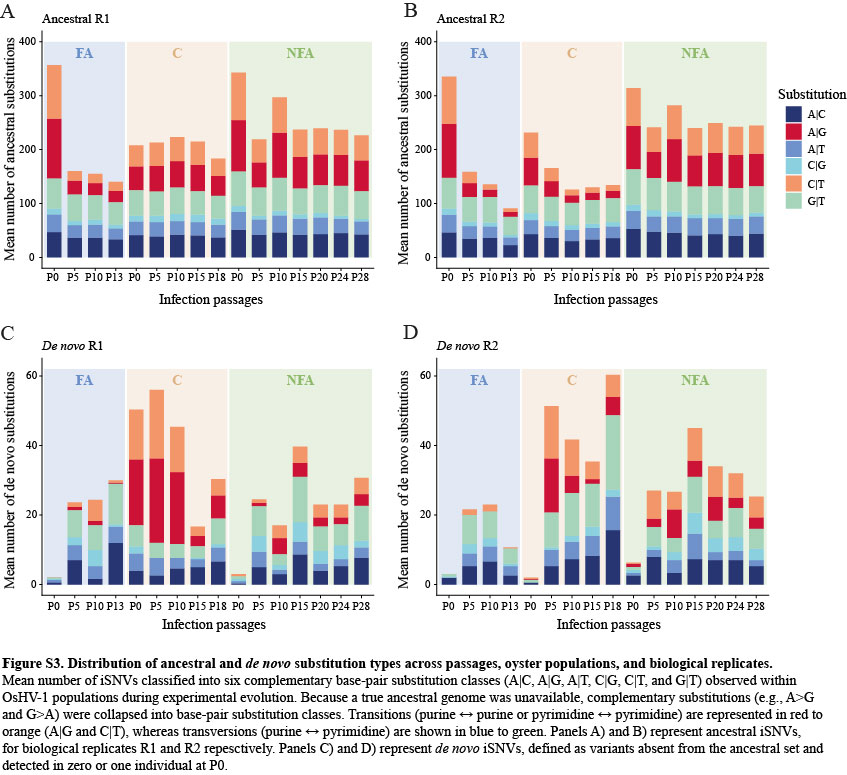

### Supplemental Figure 4

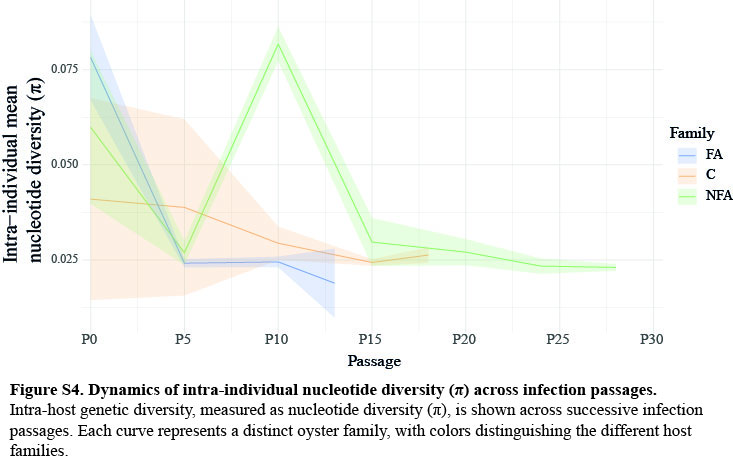
